# Dynamic Shape Modulation of Deflated and Adhered Lipid Vesicles

**DOI:** 10.1101/2025.05.06.650387

**Authors:** Gianna Wolfisberg, Jaime Agudo-Canalejo, Pablo C. Bittmann, Eric R. Dufresne, Robert W. Style, Aleksander A. Rebane

**Affiliations:** Department of Materials, ETH Zürich, 8093 Zürich, Switzerland; Department of Physics and Astronomy, University College London, London WC1E 6BT, United Kingdom; Department of Materials Science and Engineering, Cornell University, Ithaca, NY 14853, USA; Laboratory of Atomic and Solid-State Physics, Cornell University, Ithaca, NY 14853, USA; Life Molecules and Materials Lab, Programs in Chemistry and in Physics, New York University Abu Dhabi, P.O. Box 129188, Abu Dhabi, United Arab Emirates

## Abstract

Lipid membrane-bounded organelles often possess intricate morphologies with spatially varying curvatures and large membrane surface areas relative to internal volume (small reduced volumes). These features are thought to be essential for protein sorting and vesicle trafficking, but challenging to reproduce *in vitro*. Here, we show that weakly adhered giant unilamellar vesicles (GUVs) can be osmotically deflated to reduced volumes as low as 0.1, similar to what is found in flattened, disc-shaped organelles such as Golgi cisternae and ER sheets. Using shape analysis with the Canham-Helfrich model, we determine mechanical parameters including adhesion strength, membrane tension, and pressure of individual vesicles. We find that the rate of shape flattening during deflation is governed by a normalized adhesion strength that combines vesicle size, adhesion energy, and bending rigidity. For highly flattened disc-like vesicles, we identify a geometric relationship that allows the adhesion strength to be estimated solely from the vesicle’s aspect ratio, size, and bending rigidity. These results provide a quantitative experimental platform for bottom-up studies of membrane shaping mechanisms and shape-dependent phenomena, such as curvature-mediated protein sorting.

**TOC Graphic:** 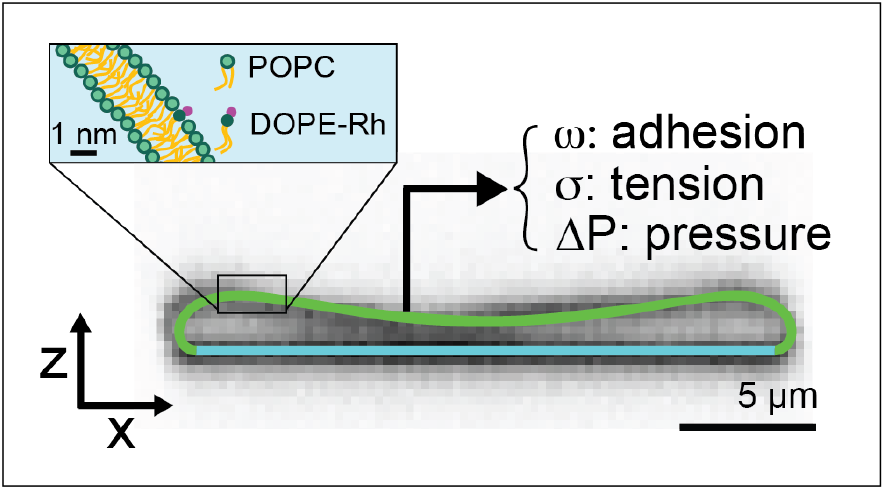

---

Eukaryotic cells contain lipid membrane-bounded compartments, each possessing distinct morphologies that facilitate specialized cellular functions. ^1–3^ While some compartments are nearly spherical, many, such as the rough endoplasmic reticulum (ER), mitochondrial cristae, chloroplast thylakoids, and Golgi cisternae, adopt disc- or sheet-like geometries characterized by extended flat regions adjoined by highly curved rims with radii of curvature in the 20 - 100 nm range.^4–10^ In the case of Golgi cisternae, the pronounced curvature difference between the flat central region and the curved rims is thought to play a critical role in the curvature-mediated sorting of cargo proteins into transport vesicles budding off from the rims.^11–14^ These structures are notable for their large membrane surface area relative to their internal volume (low reduced volume, *ν*), which provides a compact platform for protein sorting in the Golgi cisterna and maximizes the density of membrane-bound ribosomes for synthesis of secretory proteins in the rough ER.^15,16^

To elucidate the mechanisms by which flattened membrane morphologies arise and function *in vivo*, it is essential to develop model systems that replicate these shapes *in vitro* under conditions mimicking the cellular interior. Giant unilamellar vesicles (GUVs) offer a versatile platform for this purpose, particularly when their geometry and mechanical properties could be precisely controlled.^17^ Although microfluidic traps have recently been used to manipulate GUV shape, these approaches rely on physical confinement, which does not reflect the native mechanism of organelle morphogenesis.^18^ Furthermore, once fabricated, the traps offer limited control over GUV shape and curvature. Consequently, new strategies are needed to generate compartments with dynamically controlled shapes at small reduced volumes, particularly in the regime mimicking organelles such as ER sheets or Golgi cisternae, which have *ν* ≈ 0.1 or less.^8^

In general, newly produced spherical GUVs must be deflated to access the low-*ν* regime characteristic of organelle-mimetic shapes. This is typically achieved either by thermal expansion, which increases the membrane surface area while keeping the luminal volume constant,^19,20^ or osmotic deflation, which reduces the luminal volume via water efflux while keeping the membrane area fixed.^21–23^ However, these approaches have practical limitations that hinder deflation below *ν* ≈ 0.5.^24^ Thermal expansion is constrained by the small thermal area expansion coefficient of lipid bilayers, which limits achievable reduced volumes to *ν* ≈ 0.6 even at elevated temperatures of 45° C.^25,26^ Osmotic deflation, while theoretically more effective, often leads to instabilities such as pearling, tubulation, and budding transitions that arise due to Canham-Helfrich induced by asymmetry of chemical composition inside and outside the GUV.^27–34^

Even when spontaneous curvature is absent, flattened cisternal shapes remain inaccessible to free vesicles, as illustrated in Figure 1. This behavior is quantitatively captured by the Canham-Helfrich model, which describes vesicle shapes as a result of minimizing bending energy subject to constraints on the surface area and volume. ^35,36^ At reduced volumes below *ν* ≲ 0.6, the model predicts that free vesicles adopt curved, cup-like configurations (stomatocytes), rather than fattened discs. ^37–40^ Experimental observations have confirmed these predictions.^38,41^ However, adhesion to a flat substrate such as a microscope coverslip or a supported lipid bilayer (SLB) can suppress the formation of stomatocytes and instead stabilize flattened shapes. ^42,43^ Under these conditions, the global vesicle shape is governed by a balance between reduced volume *ν* and the adhesion energy per unit area, *ω*. In the strong adhesion limit, vesicles approximate spherical caps with a contact angle determined by *ν*, as bending energy is negligible and the vesicle tries to maximize its adhesion area against membrane tension, similar to a liquid droplet wetting a surface. ^44,45^ At intermediate adhesion strengths, GUVs can adopt a variety of non-spherical morphologies with distinct curvature profiles outside the adhesion zone. ^25,26,46^ These regimes have been experimentally achieved using non-specific adhesion such as depletion interaction, van der Waals interaction, or an applied electric field. ^25,26,46–49^ Nonetheless, achieving and maintaining stable, flattened shapes at reduced volumes below *ν* ≈ 0.5 has remained challenging.

**Figure 1:**
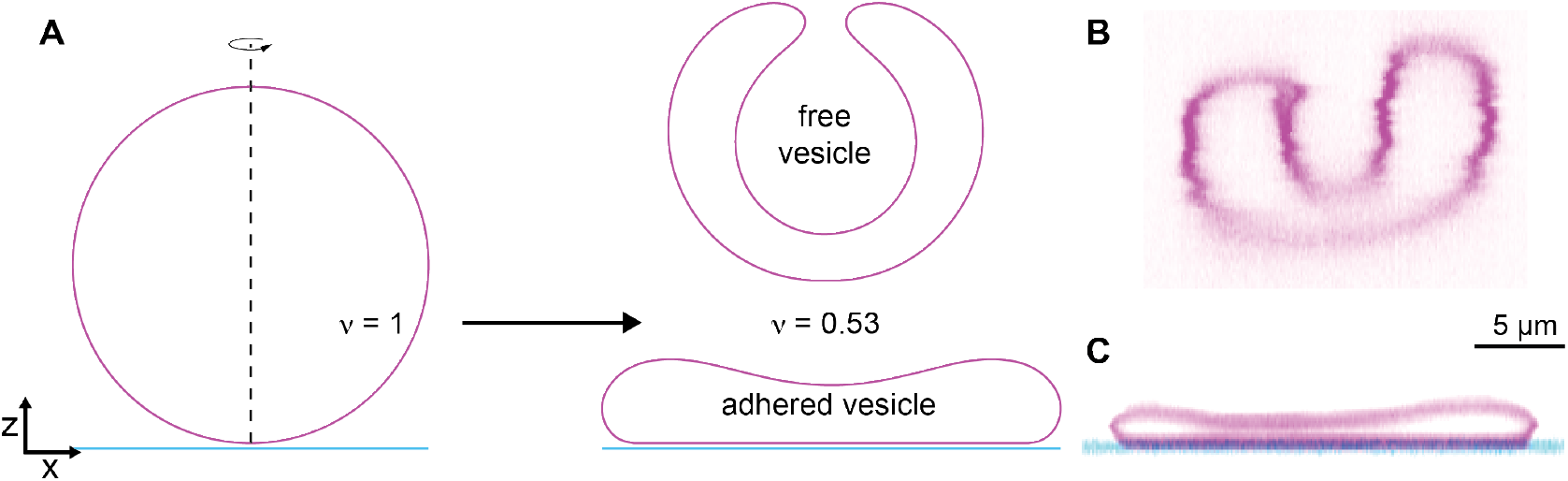
Adhesion flattens deflated vesicles. **(A)** Shape transformations calculated using the Canham-Helfrich model from fully inflated (reduced volume *ν* = 1) to deflated (*ν* = 0.53). The free vesicle (top) curls up into a cup-like stomatocyte whereas the adhered vesicle can assume a flat shape. **(B)** Corresponding confocal fluorescence micrographs of a free vesicle with *ν* = 0.69 and **(C)** an adhered vesicle with a reduced volume of *ν* = 0.26. Note that the free vesicle shown here has a reduced volume above the stomatocyte transition prediced by the Canham-Helfrich model, likely due to the effect of flows on the free vesicle.

Here, we present an approach to stably deflate adhered GUVs down to *ν* ≈ 0.1 and characterize their morphologies using a Canham-Helfrich model. We show that weak adhesion is essential to stabilize vesicles at such low reduced volumes and prevent the onset of shape instabilities. By tuning the osmotic pressure and adhesion strength, we observe a smooth, controllable progression of vesicle morphologies from spherical caps to flattened discocytes. From the deflation trajectories, we find that the rate of shape flattening is governed by a normalized adhesion strength that incorporates vesicle size, adhesion energy, and bending rigidity. Furthermore, for flat, disc-like vesicles, we identify a predictive relationship that allows the adhesion strength to be estimated solely from the vesicle’s aspect ratio, size, and bending rigidity, providing a geometric handle on key mechanical parameters without the need for numerical shape fitting.

## Results and Discussion

### Experimental Design

In order to observe individual GUVs during successive steps of osmotic deflation, we constructed a diffusion chamber (Fig. S8). Briefly, we created a passive flat substrate for the GUVs by coating a microscope cover glass with a SLB of 1-palmitoyl-2-oleoyl-glycero-3-phosphocholine (POPC) (Fig. 2A). We made a sample chamber by sticking a thin (120 µm) annular spacer on the bilayer and filled it with electroformed GUVs (25 mM sucrose in the lumen, 99.9 mol% POPC and 0.01 mol% 1,2-dioleoyl-*sn*-glycero-3-phosphoethanolamine-N-(lissamine rhodamine B sulfonyl) (Rh-DOPE)) suspended in an isotonic solution (25 mOsm/L) containing 10 mM NaCl, 5 mM glucose, and 0.2, 0.4, or 0.8 % (w/v) PEG 100 kDa. The presence of PEG induced weak tunable adhesion between the GUV and the SLB via the depletion interaction.^50^

**Figure 2:**
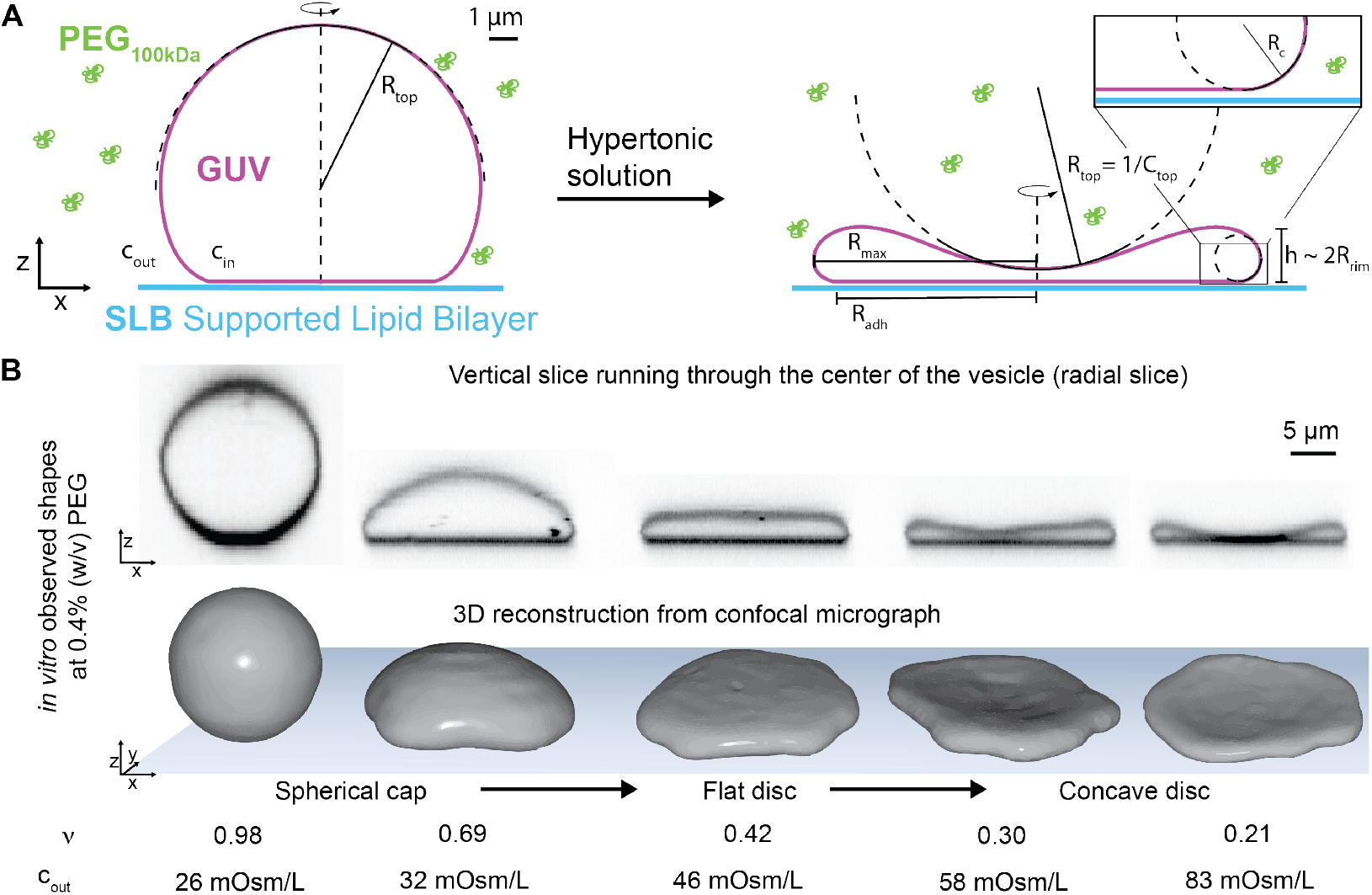
Shape transformations accompanying deflation of adhered giant unilamellar vesicles (GUVs). **(A)** Schematic of a GUV adhered to a supported lipid bilayer (SLB) before (left) and after (right) deflation via the addition of a hypertonic solution. The presence of PEG 100 kDa (*green*) induces adhesion between the GUV and SLB via the depletion interaction. The concentration of PEG remains constant throughout a sequence of shape transformations. *c*_*in*_ and *c*_*out*_ indicate osmolarity inside and outside of the GUV, respectively. *R*_*top*_ indicates the signed radius of curvature of the top of the vesicle with *R*_*top*_ *>* 0 for convex and *R*_*top*_ *<* 0 for concave vesicles and *C*_*top*_ = 1*/R*_*top*_ is the curvature at the top of the vesicle. **(B)** Top: Color-inverted confocal images of orthogonal cross-sections (xz) showing a typical sequence of shape transformations of a single GUV in the presence of 0.4% (w/v) PEG 100 kDa and a gradual increase of external osmolarity *c*_*out*_. Bottom: Corresponding 3D-reconstructions of the GUVs. The GUV shape transitions from a convex adhered spherical cap to a flat adhered disc and finally to a concave adhered disc. The corresponding reduced volume 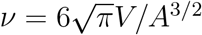 is given under each shape.

As the GUVs sedimented to the bottom of the chamber, we filled a disposable dialysis cup (3.5 kDa molecular weight cutoff) with deflation buffer and placed it on top of the spacer, bringing the dialysis membrane in direct contact with the GUV solution. By incrementally increasing the osmolarity via the glucose concentration of the deflation buffer, we could gradually deflate the GUVs to the desired *ν* while maintaining a constant concentration of the depletant PEG. The dialysis membrane effectively suppressed any flows arising from glucose density gradients, allowing us to keep track of individual GUVs over the course of hours and successive exchanges of deflation buffer. The system reached equilibrium within approximately 1 hour at each deflation step, consistent with the time-scale of achieving a homogeneous glucose distribution throughout the sample chamber.

### Controlled Deflation of Adhered GUVs

Our diffusion chamber allowed us to observe a striking sequence of GUV shape transitions accompanying deflation across a broad range of reduced volumes. A typical deflation series of a single GUV in the presence of 0.4% (w/v) PEG is shown in Fig. 2B. We acquired approximately 60 deflation series at three adhesion strengths corresponding to 0.2%, 0.4%, and 0.8% (w/v) PEG. The GUV and SLB membranes maintained contact without lipid mixing, indicating that the bilayers remained stable and did not undergo full fusion or hemifusion (Fig. S1). In our subsequent analysis, we excluded vesicles that harbored significant defects such as tubular structures, resulting in a data set of N=21 vesicles across the three PEG concentrations. We generated 3D-reconstructions of the membrane surfaces (Fig. 2B) and calculated the GUV volume, *V*, surface area, *A*, reduced volume 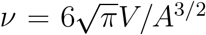 and adhered surface area, *A*_*adh*_, at each stage of deflation. Notably, we found that these observables could be obtained from any vertical slice passing through the center of a vesicle (radial slice) and, by assuming axisymmetry, yielded similar values to those obtained from the significantly more cumbersome reconstruction of the full three-dimensional shapes (Fig. S2).

Osmotic deflation relies on exposing the GUV with internal osmolarity *c*_0_ to a hypertonic solution with osmolarity *c*_*out*_ *> c*_0_. This induces water efflux across the lipid membrane until osmotic equilibrium is reached, *i*.*e. c*_*in*_ = *c*_*out*_ within Δ*c* ≈ 1 mOsm/L.^51^ We assume that the GUV lumen behaves as an ideal solution following van ‘t Hoff’s equation and *c*_*in*_ = *n/V*, where *V* is the GUV volume and *n* is the number of sucrose molecules inside the GUV. The bilayer is impermeable to sucrose and *n* remains approximately constant during deflation, *n* = *V c*_*in*_ = *V*_0_*c*_0_, yielding the equilibrium condition for the reduced volume

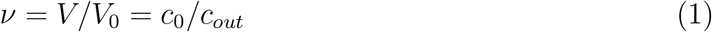

for a vesicle that is spherical at osmolarity *c*_0_.

Indeed, we observed that the reduced volumes obey Eq. 1 with *c*_0_ = 25 mOsm/L (Fig. 3A), confirming that the GUVs have reached osmotic equilibrium. We found that the deflation is reversible (Fig. 3B), demonstrating that our setup allows us to control the reduced volume via osmolarity of the deflation buffer. In some cases, the adhesion area did not fully retract upon re-inflation, possibly due to some pinning of the contact line. The GUV surface area remained constant over the course of the entire deflation series (Fig. 3C), consistent with the incompressibility of lipid bilayers. It is therefore convenient to adopt the radius of the fully inflated GUV, 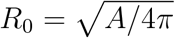, as a length scale.

**Figure 3:**
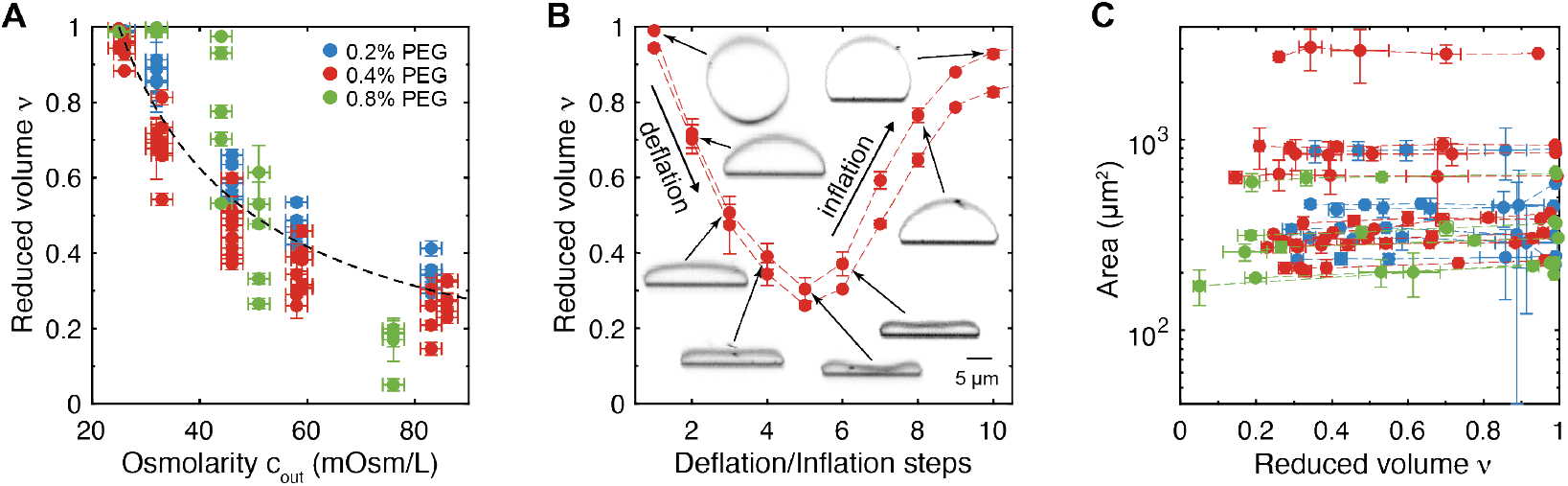
Osmotic deflation reversibly changes the reduced volume while the surface area remains constant. **(A)** Measurements of the GUV’s reduced volume at increasing outside osmolarities *c*_*out*_ at three PEG concentrations: 0.2% (w/v) (*blue*), 0.4% (w/v) (*red*) and 0.8% (w/v) (*green*). The black dashed line shows the expected values for osmotic equilibrium, *ν* = *c*_0_*/c*_*out*_ with *c*_0_ = 25 mOsm/L. **(B)** Reduced volumes of two typical GUVs undergoing deflation and subsequent inflation following changes in *c*_*out*_. **(C)** Total GUV area measured at different deflation steps as a function of the reduced volume. Data points belonging to the same GUV are connected by a dashed line. Error bars in (A) - (C) are standard deviations among measurements obtained from 3 different confocal slices at 120° angles, except for the osmolarity in (A), in which an uncertainty of 2 mOsm/L was assumed based on the precision of the osmometer.

### Shape Evolution During Deflation

At all three adhesion strengths, deflation was accompanied by a sequence of shape transformations starting with a convex adhered spherical cap, followed by a flattened disc, and ending with a concave adhered discocyte as shown in Fig. 2B for 0.4% PEG (see Figs. S3 - S5 for deflation series at each of the three PEG concentrations). Under isotonic conditions (25 mOsm/L) the GUVs were nearly spherical or adhered to the SLB while closely approximating the shape of a spherical cap with an effective contact angle *θ >* 90° and an adhered surface area that was constrained by the fixed GUV volume and total surface area (Fig. 2B, *ν* = 0.98). The first deflation step (*ν* = 0.69) allowed the adhered area to expand while reducing the contact angle. At the second step (*ν* = 0.42), the vesicles adopted a disc-like shape with a flat top and rounded rims that had no well-defined effective contact angle. In the third step (*ν* = 0.30), the shape became an adhered discocyte with a concave top, where the rims maintained a curvature similar to that of the flattened disc. Further deflation (*ν* = 0.21) bent the top of the adhered discocyte inward until it was forced to flatten due to contact with the adhered membrane at the opposite end of the GUV.

In the limit of very strong adhesion the bending rigidity of the membrane becomes negligible and the adhered GUVs form perfect spherical caps with effective contact angles that are exactly determined by the reduced volume via

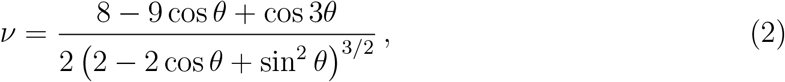

independent of bending rigidity and adhesion strength (Fig. S6, *black dashed*).^44^ The effective contact angles of deflated vesicles in the presence of 0.4% and 0.8% PEG were consistently lower than those of perfect spherical caps at the same reduced volumes (Eq. 2), revealing the influence of bending rigidity on the vesicle geometry in the strong adhesion regime (Fig. S6, *symbols*). Notably, vesicles in 0.2% PEG did not form spherical caps with well-defined contact angles even for high reduced volumes, suggesting that bending rigidity significantly counteracted vesicle deformation due to adhesion. The degree to which deflation affects the shape of a vesicle depends on the adhesion strength, the bending rigidity, and the vesicle size. The change in GUV shape accompanying deflation can be characterized by the signed radius of curvature at the top of the vesicle, *R*_*top*_ (Fig. 2A). The normalized top curvature, *Ĉ*_*top*_ = *R*_0_*/R*_*top*_, decreased with *ν* (Fig. 4A and B) where *Ĉ*_*top*_ *>* 0 indicates spherical cap-like geometries, *Ĉ*_*top*_ = 0 flattened disc-like shapes, and *Ĉ*_*top*_ *<* 0 adhered discocytes.

**Figure 4:**
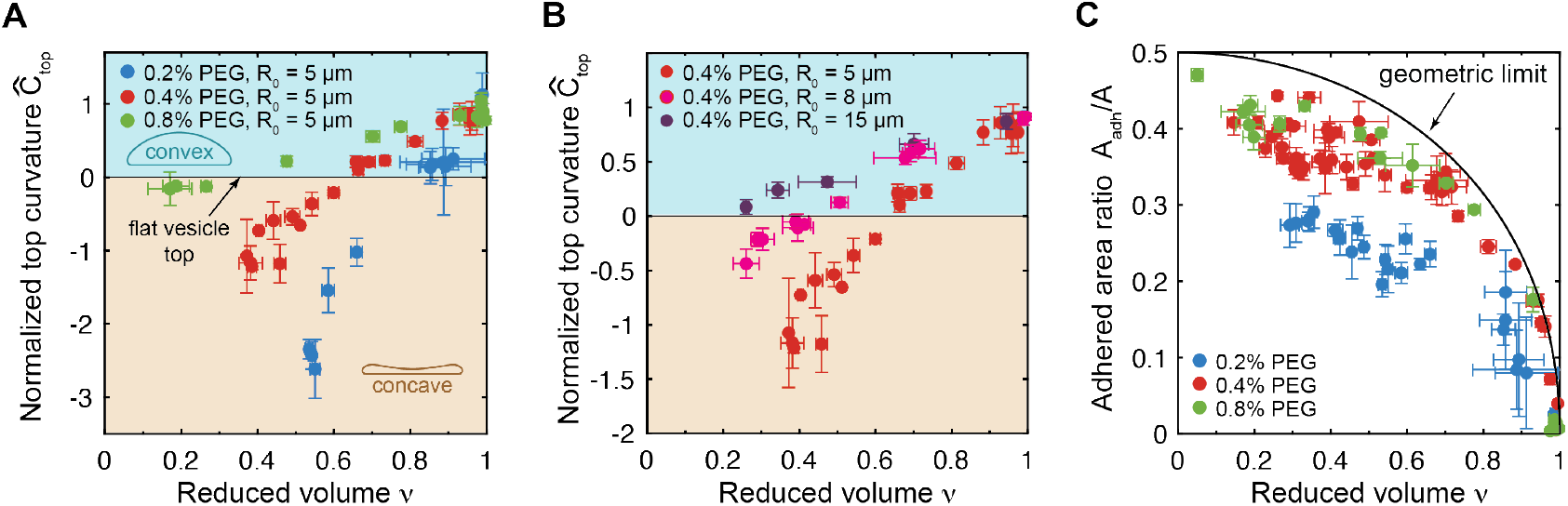
Changes in the top curvature and adhered area of GUVs accompanying deflation. **(A)** Normalized top curvatures, *Ĉ*_*top*_ = *R*_0_*/R*_*top*_, as a function of reduced volume for different PEG concentrations (adhesion strengths) for vesicles of similar size (*R*_0_ = 5 ± 1 *µ*m). **(B)** Normalized top curvature as a function of the reduced volume at the same PEG concentration of 0.4% (w/v) but for vesicles of different sizes, *R*_0_ = 5±1 *µ*m (*red*), *R*_0_ = 8±1 *µ*m (*magenta*), and *R*_0_ = 15 ± 1 *µ*m (*purple*). **(C)** Ratio of adhered vesicle surface area, *A*_*adh*_, to total vesicle surface area, *A*, as a function of reduced volume at different PEG concentrations. The adhered area ratio in the limit of infinitely strong adhesion (geometric limit) is shown by the black curve. Colors in (A) and (D) indicate the concentration of PEG 100 kDa in the outside buffer: 0.2% (w/v) (*blue*), 0.4% (w/v) (*red*), 0.8% (w/v) (*green*). Error bars in (A)-(D) and (E) are standard deviations among measurements obtained from 3 different confocal slices at 120° angles.

Comparison of vesicles of similar size (*R*_0_ ≈ 5 *µ*m) but at different concentrations of PEG reveals that the rate of change of *Ĉ*_*top*_ as a function of *ν* depends on the adhesion strength (Fig. 4A), whereby vesicles in the presence of stronger adhesion (*green*) flatten more slowly with deflation than those in the presence of weaker adhesion (*blue*). Inflated spherical caps attain the greatest positive top curvature approaching 1*/R*_0_ for *ν* ≈ 1, regardless of the adhesion strength. The greatest negative top curvature was achieved for adhered discocytes with the weakest adhesion (0.2% PEG). In the presence of 0.2% PEG, weakly adhered flattened disc- or cigar-like shapes were already achieved at the first deflation step (Fig. S3). The cigar-like shapes occurred only at around *ν* ≈ 0.9 and were not axisymmetric, resulting in large error bars for their reduced volume, *Ĉ*_*top*_, and adhered area fraction (Fig. S3A and C). However, further deflation restored axisymmetry by transforming the cigar-like shapes into adhered discocytes. A similar comparison for vesicles at the same PEG concentration (0.4% PEG) but different *R*_0_ shows that the top curvature changes more quickly with deflation for smaller vesicles (Fig. 4B). The broadest range of *C*_*top*_ = 1*/R*_*top*_ was achieved for vesicles with *R*_0_ = 5 *µ*m in the presence of 0.2% PEG, allowing us to control the top curvature between *C*_*top*_ = −5 *µ*m^*−*1^ and *C*_*top*_ = +0.2 *µ*m^*−*1^ by controlling the osmolarity outside the GUV.

The adhered surface area fraction *A*_*adh*_*/A* increased monotonically with increasing deflation (Fig. 4C). In the limit of small reduced volumes and in the presence of 0.4% or 0.8% PEG, the adhered area fraction approached 0.5, which corresponds to a thin pancake that is adhered to the membrane on one side. Vesicles in the presence of 0.2% PEG were generally more dynamic and tended to detach from the membrane at small reduced volumes, forming biconcave discocytes with no well-defined adhered area fraction (Fig. S3). Adhesion leads the vesicle to maximize its adhered surface area until a balance between adhesion strength, membrane tension, and bending elasticity is achieved. In the strong adhesion limit expansion of the adhered surface area is limited by membrane tension. For perfectly axisymmetric vesicles, *A*_*adh*_*/A* then only depends on the reduced volume via

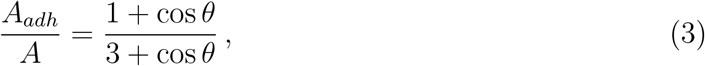

where *θ* is implicitly defined by the reduced volume using Eq. 2 (Fig. 4C, *black*). Deflation lowers the membrane tension, which immediately is compensated for by an increase in the adhered surface area. In the presence of finite adhesion, however, the adhered area is additionally limited by the contribution of membrane bending rigidity. The first order approximation for *A*_*adh*_*/A* for adhered axisymmetric vesicles in the presence of bending rigidity is a function of 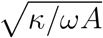 and the reduced volume *ν*:^44^

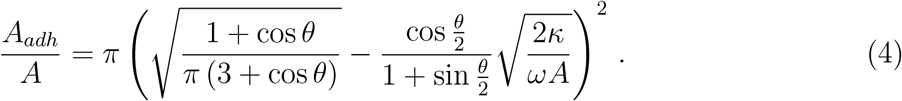

For slightly deflated adhered vesicles whose shapes closely approximate spherical caps with well-defined contact angles, it is possible to estimate the adhesion strength based on the vesicle reduced volume and the adhered area fraction. ^44,49^ In our case of highly deflated vesicles that do not have a well-defined contact angle, however, this approximation breaks down.

### Mechanical Parameters from Shape Analysis

To extract the mechanical parameters of GUV membranes, including adhesion strength *ω*, membrane tension *σ*, and pressure difference Δ*P* = *P*_*in*_ − *P*_*out*_, we took advantage of the axisymmetry of the vesicles in our dataset. We wrote a custom Matlab script that compares a radial slice (vertical confocal slice running through the center of a vesicle) to shapes obtained from numerical integration of the axisymmetric shape equations assuming zero spontaneous curvature (see Materials and Methods in Supporting Information for details). Figure 5A-C shows overlays of the numerical calculations and the experimental observations of the vesicle shown in Fig. 2B, which were determined by manual adjustment until good agreement between the two was achieved. The mechanical parameters corresponding to each shape were converted from dimensionless values assuming a membrane bending rigidity *κ* = 33 k_B_T for POPC membranes, where *k*_*B*_*T* is the Boltzmann constant and *T* is room temperature.^52^ The calculated shapes fit the observations well, despite the deviations from perfect axisymmetry that can be seen in the 3D-reconstructions shown in Fig. 2B. Extraction of mechanical parameters of GUV membranes has previously been achieved by using using Surface Evolver to model three-dimensional membrane shapes observed with confocal microscopy.^24–26^ Although this approach has the advantage of being applicable to non-axisymmetric vesicles, it requires full three-dimensional reconstruction and modeling of the membranes shapes, which can be challenging and time-intensive to use for nonexperts. In contrast, the approach we present here requires at minimum only one radial slice of an approximately axisymmetric vesicle to obtain the desired physical quantities.

**Figure 5:**
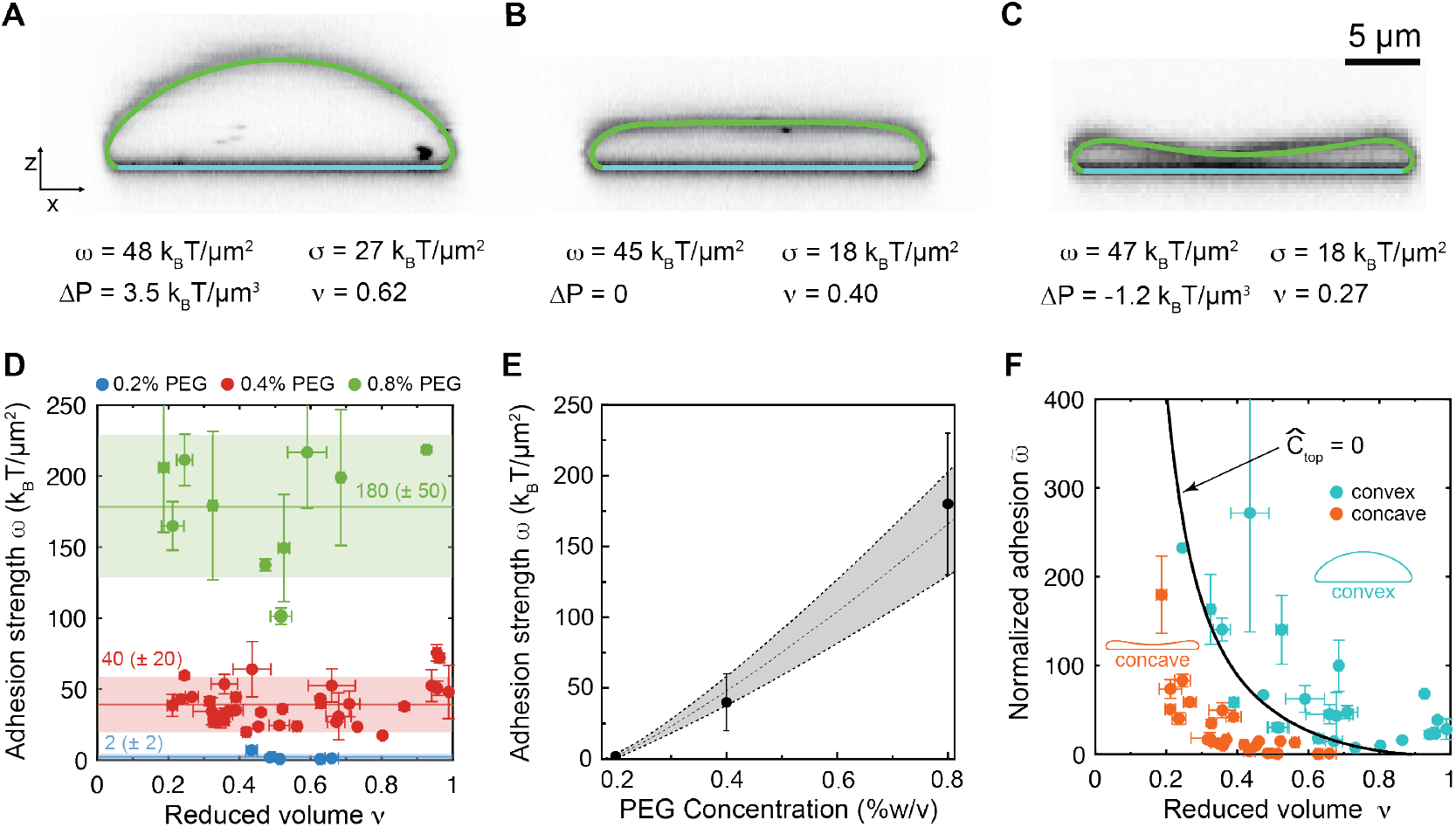
Quantification of adhesion strength through shape analysis. **(A-C)** Overlays of numerically calculated shapes from the Canham-Helfrich model on vesicles observed with confocal fluorescence microscopy. Colors represent non-adhered membrane (*green*) and adhered membrane (*cyan*). Tabulated are the adhesion strength *ω*, membrane tension *σ*, pressure difference Δ*P*, and reduced volume *ν* corresponding to each shape, assuming a bending rigidity of *κ* = 33 k_B_T. The non-dimensionalized best-fit model parameters for each shape along with the complete set of observables are given in Table S1. **(D)** Adhesion strengths determined by shape-fitting for three different concentrations of PEG 100 kDa: 0.2% (w/v) (*blue*), 0.4% (w/v) (*red*), 0.8% (w/v) (*green*). Typical fits are shown in (A)-(C). The horizontal lines and shaded regions show the mean and standard deviations, respectively, of measurements obtained from all shapes at a particular PEG concentration. **(E)** Measured adhesion strength as a function of PEG concentration. The dashed curve and shaded area represent the best-fit curve and its 68% confidence interval, respectively, using the model for depletion strength described by Eq. 7 with best-fit parameters *ξ* = 0.04 ± 0.03 and *l* = 9 ± 2 nm. The means and standard deviations of the plotted adhesion strengths correspond to the values quoted in (D). **(F)** Normalized adhesion strength 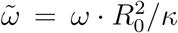 as a function of reduced volume. Colors denote convex (*R*_*top*_*/R*_0_ *>* 0, *cyan*) and concave (*R*_*top*_*/R*_0_ *<* 0, *orange*) vesicles. The black curve represents solutions from the Canham-Helfrich model with *C*_*top*_ = 0 and assuming zero spontaneous curvature, *C*_0_ = 0. Error bars in (D) and (F) are standard deviations among measurements obtained from 3 different confocal slices at 120° angles.

We determined the adhesion strength from the fitted shape using 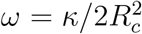, where *R*_*c*_ is the radius of curvature of the fitted shape along the contour running from top to bottom of the vesicle, at the point where the vesicle meets the SLB (see illustration in Fig. 2A).^42^ As expected, the adhesion strength remained approximately constant during deflation at constant PEG concentration (0.4% PEG).

The vesicle starts with an overpressure for the convex shape at (Fig. 5A, *ν* = 0.62), reaches zero for the flat, disc-like shape at (Fig. 5B, *ν* = 0.40), and transitions into an underpressure for the adhered discocyte at (Fig. 5C, *ν* = 0.27), reminiscent of the change in Laplace pressure across a purely tension-dominated interface (*e*.*g*. a liquid droplet) as the interface’s curvature changes sign. We find that a pressure difference of 3.5 k_B_T*/µ*m^3^ corresponds to a negligible osmolarity difference of Δ*c* = 6 nM ≪ *c*_*in*_, confirming that the externally applied osmolarity of the deflation buffer controls the reduced volume.

The membrane tension initially decreases with deflation as the vesicle transitions from a convex to a flat, disc-like shape, and remains constant for further deflation beyond the disc-like shape (Fig. S7). The average membrane tension of the three stages of deflation is *σ* ≈ 23 ±5 k_B_T*/µ*m^2^, which is small compared to the membrane lysis tension of *σ*_*lys*_ ≈ 4 mN*/*m = 10^6^ k_B_T*/µ*m^2^.^53^ Bending energy dominates the vesicle shape in the highly curved regions near the contact line and the vesicle rims, but at the top of the vesicle its contribution is reduced, particularly for flat vesicles. We can define the bendocapillary length, 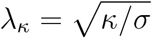, to assess the relative contributions of bending and membrane tension to the mechanics of the top of the vesicle.^54^,55 The top of the vesicle is dominated by membrane tension for *λ*_*κ*_ ≪ *R*_*top*_, and by bending for *λ*_*κ*_ ≫ *R*_*top*_. For the vesicle shown in Fig. 5A-C, we get *λ*_*κ*_ ≈ 1.2 ± 0.1 *µm* ≪ *R*_*top*_, indicating that the bending energy is negligible at the top of the vesicle. In other words, the curvature is determined by membrane tension, similar to a liquid droplet. We therefore asked whether the membrane curvature, tension, and pressure are related by the Young-Laplace equation,

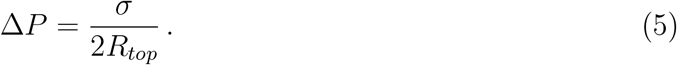

Using the values from Table S1 for the vesicle shown in Fig. 5A-C we found good agreement with Eq. 5 for all three stages of deflation. Axisymmetric shape analysis therefore offers insight into which mechanical parameters are decisive for local vesicle geometries.

### Tunable Adhesion Using Depletion Interactions

The depletion-induced adhesion between two flat surfaces is well described by the Asakura-Oosawa-Vrij (AOV) model assuming ideal solution behavior and a perfectly non-adsorbing depletant:

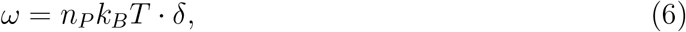

where *n*_*P*_ is the number density of depletant (PEG) and *δ* is the thickness of the depletion layer on the surfaces.^56–58^ Treating PEG as a polymer in good solvent yields *δ* ≈ *R*_*g*_, where *R*_*g*_ is the radius of gyration of PEG.^59^ However, whereas the SLB is flat over the relevant length scales, GUV membranes with low tensions undergo thermal undulations that effectively reduce the thickness of the depletion layer:

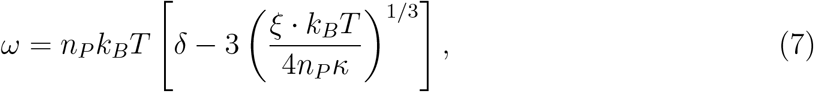

where *ξ* is a non-dimensional parameter quantifying the undulations, predicted to lie in the range 0.01-0.23.^52,60,61^

Using shape analysis, we extracted the adhesion strengths for all vesicles at different stages of deflation and at different PEG concentrations (Fig. 5D). Increasing the PEG concentration increased the adhesion strength, in agreement with the concentration-dependence of the depletion interaction. Averaging over our entire dataset yielded *ω* = 180±50 *k*_*B*_*T/µ*m^2^ for 0.8% PEG, *ω* = 40 ± 20 *k*_*B*_*T/µ*m^2^ for 0.4% PEG, and *ω* = 2 ± 2 *k*_*B*_*T/µ*m^2^ for 0.2% PEG. We fitted this concentration-dependence with Eq. 7, yielding best-fit parameters with *ξ* = 0.04 ± 0.03 and a depletion thickness of *δ* = 9 ± 2 nm (Fig. 5E). Both parameters fall into the expected range, however, the depletion thickness of *δ* is somewhat lower than the radius of gyration of PEG 100 kDa (*R*_*g*_ ≈ 16 nm), indicating that PEG does not act as a perfectly non-adsorbing depletant.^62^ Indeed, molecular dynamics simulations have predicted weak binding of PEG to lipid bilayers membranes with interaction energies on the order of 1 k_B_T per PEG molecule.^63^ Thus, vesicle shape analysis allows us to determine GUV adhesion strengths that agree with quantitative models of the depletion interaction.

Next, we asked how strongly the shape of each adhered GUV responded to deflation. We quantified this with the reduced volume at which each vesicles adopts a flat, disc-like shape. Because this reduced volume depended on both the adhesion strength and the vesicle size (Fig. 4A and B), we postulated that it would be fully determined by a normalized adhesion strength that combines these two parameters, 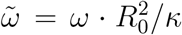. We plotted the normalized adhesion strength against *ν* and labeled the points as convex (Fig. 5F, *cyan*) or concave (*orange*) based on the sign of their top curvatures. The plot reveals a neat segregation of concave and convex vesicles into two areas that are separated by a border marking flat disc-like vesicles with vanishing top curvature. To obtain a theoretical prediction for this border, we numerically solved the shape equations for adhered axisymmetric vesicles by setting *Ĉ*_*top*_ = 0, yielding a curve that overlaps with the border between experimentally observed shapes of convex and concave vesicles (Fig. 5F, *black*). Thus, given a vesicle size *R*_0_, adhesion strength *ω*, and bending rigidity *κ*, we could accurately predict the degree of deflation needed to shape adhered GUVs into flat disc-like shapes.

### Flat Adhered Disc-like Vesicles

Vesicles with nearly vanishing top curvature form an important class of shapes found in subcellular organelles such as Golgi cisternae. To investigate the properties of these shapes, we chose to look at vesicles with normalized top curvature −0.25 *< Ĉ*_*top*_ *<* 0.25 (Fig. 6A). These shapes closely resembled thin pancakes, whose total area is the sum of two discs of radius *R*_*max*_, where *R*_*max*_ is the vesicle’s radius at its widest point. It is therefore possible to accurately estimate *R*_0_ and hence the total vesicle area just by measuring *R*_*max*_ within a radial slice of a disc-like vesicle (Fig. 6B). An important characteristic of these shapes is the high curvature region at the rim of the disc. While the curvature varies smoothly from ≈ 0 at the top of the vesicle to 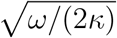 at the point of contact with the substrate, ^42^ a characteristic radius of curvature of the rim can be defined as being half the height *h* of the vesicle, *R*_*rim*_ = *h/*2 (Fig. 2A). Interestingly, we found that the normalized radius of curvature at the rim decreased with reduced volume in a manner that is independent of the adhesion strength (Fig. 6C). This agrees well with numerical solutions of the shape equations where we defined disc-like shapes using the boundary condition *Ĉ*_*top*_ = 0 (Fig. 6C, *black*).

**Figure 6:**
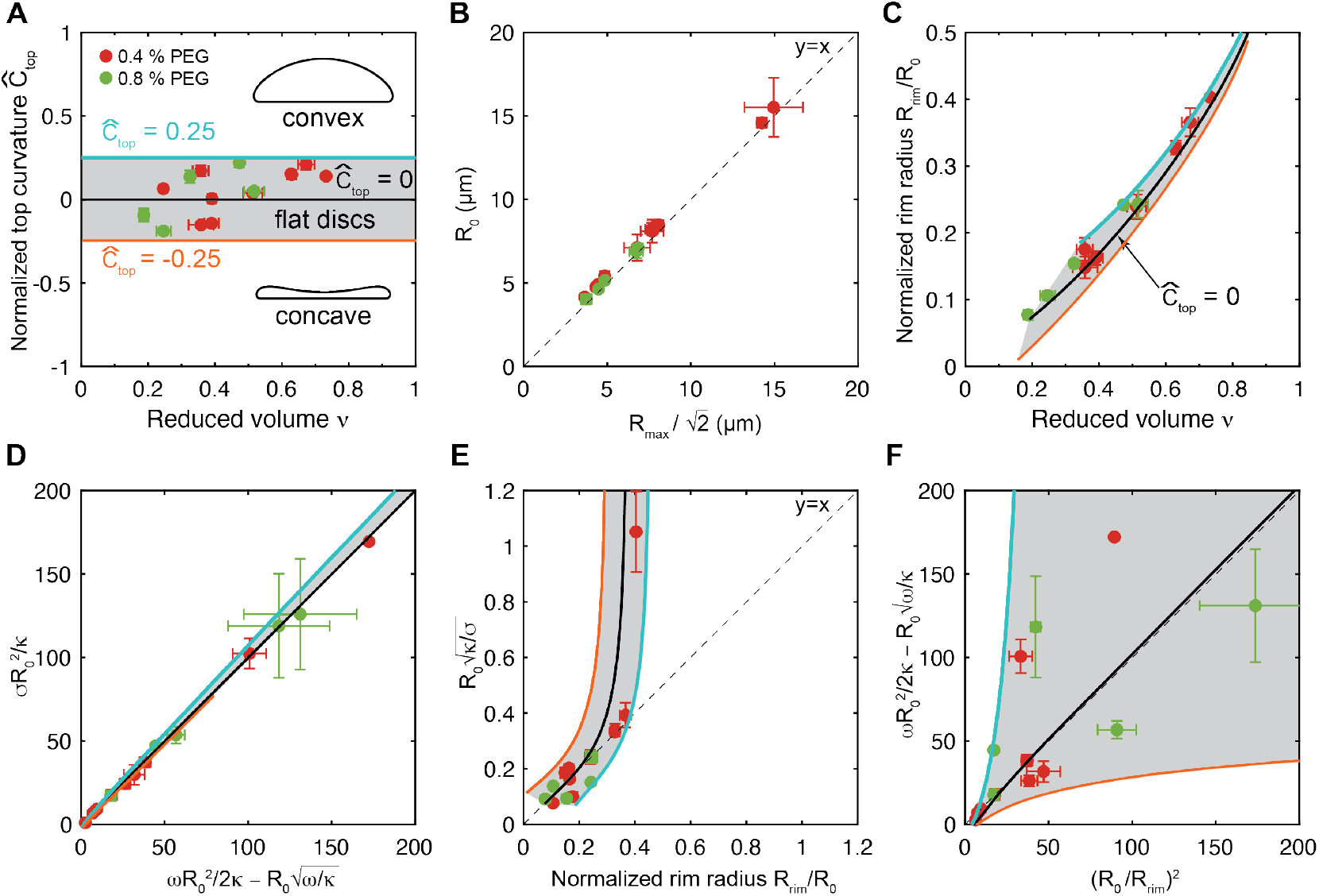
Geometrical and mechanical properties of flat adhered disc-like vesicles. **(A)** Flat adhered disc-like vesicles in (A) - (F) are defined as those having a normalized top curvature (*Ĉ*_*top*_ = *R*_0_*/R*_*top*_) within −0.25 *< Ĉ*_*top*_ *<* 0.25, denoted by the gray shaded area. **(B)** Vesicle size *R*_0_ as a function of the scaled maximum disc radius 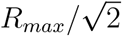. **(C)** Normalized rim curvature, which coincides with the aspect ratio of the vesicle, as a function of the reduced volume. **(D)** Normalized membrane tension versus the normalized prediction from modified Young’s law.^44,64^ **(E)** Normalized bendo-capillary length as a function of the normalized rim radius, *R*_*rim*_, defined as half the vesicle height as shown in Fig. 2A. **(F)** Universal relationship between aspect ratio *R*_0_*/R*_*rim*_ and the adhesion strength, vesicle size, and bending rigidity for adhered vesicles with flat tops. Black solid curves in (A) and (C) - (F) represent solutions from the Canham-Helfrich model, with top curvature bounds *Ĉ*_*top*_ = 0.25 (*blue*) and *Ĉ*_*top*_ = − 0.25 (*orange*). Colors indicate the concentration of PEG 100 kDa in the outside buffer: 0.4% (w/v) (*red*) and 0.8% (w/v) (*green*). Gray dashed lines indicate equal ratio of x and y. Confocal microscopy images of all flat disc-like vesicles can be found in Fig. S10.

In the limit of strong adhesion, adhered vesicles form spherical caps whose membrane tension is expected to follow Young’s law, *σ* = *ω/*(1 + cos *θ*), where *θ* is the effective contact angle.^44^ In the presence of finite adhesion, however, this expression must be modified to account for the effect of membrane bending, yielding

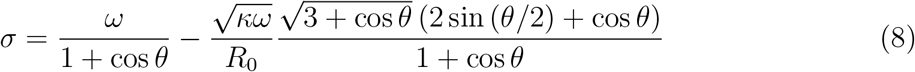

for approximately spherical caps, where the second term accounts for the contribution of bending to the vesicle shape. ^49^ For disc-like vesicles, the contact angle approaches *θ* = 0 and Eq. 8 simply becomes

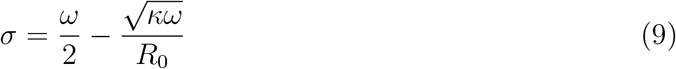

We found that the membrane tension and adhesion strength of disc-like vesicles closely followed Eq. 9 (Fig. S11A), which is somewhat surprising given the apparent differences between these shapes and the spherical cap geometry. Eq. 9 reveals that the tension is independent of the reduced volume (Fig. S7) and is instead primarily determined by the adhesion strength and, secondarily, by the bending rigidity and the vesicle size. For small normalized rim radii *R*_*rim*_*/R*_0_ *<* 0.3, the vesicle rim is well approximated by a syntractrix shape that balances the curvature elasticity with the membrane tension, ^54^ with *R*_*rim*_ ≈ *λ*_*κ*_ (Fig. 6E, see Materials and Methods in Supporting Information for details). This approximation holds whenever the principal radius of curvature at the rim along the contour from top to bottom of the vesicle (of order ~ 1*/R*_*rim*_) is much greater than the radius along the equator of the vesicle (of order ~ 1*/R*_0_). If this condition is satisfied, the height of disc-shaped vesicles, *h* = 2 · *R*_*rim*_, is determined by the surface tension and bending rigidity, independent of vesicle size or adhesion strength (Fig. S11B). By combining the previous two results, we find a universal relationship between the adhesion strength, vesicle size, and the aspect ratio of disc-like vesicles (Fig. 6F, *black dashed*),

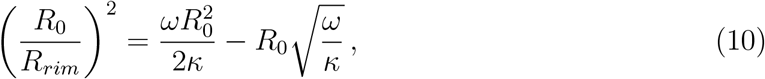

which closely approximates the exact numerical solution of the shape equations for *C*_*top*_ = 0 (Fig. 6F, *black solid*). In other words, using Eq. 10, it is possible to determine the adhesion strength of flat adhered vesicles based on their size, rim height, and the bending rigidity. However, the accuracy of this relationship rapidly decreases with decreasing normalized rim size and increasing normalized adhesion strength.

### Rim Curvature of Highly Deflated Vesicles

Deflation beyond the flattened discs yielded adhered discocytes with concave tops. The concave top bent inward and eventually flattened out as it made contact with the opposite, SLB-bound GUV membrane (Fig. 7). We did not solve the shape equations with this boundary condition and therefore could not extract the mechanical parameters of this family of shapes. However, we found that the rim curvatures appeared to steadily increase with decreasing reduced volume, falling below the bendo-capillary length. The highest curvatures were achieved for the smallest vesicles and for the highest adhesion strengths (Fig. 7). The adhesion experienced by vesicles in the presence of 0.2% PEG (2 k_B_T*/µ*m^2^) was strongly counteracted by bending such that for reduced volumes below *ν* ≈ 0.3, the vesicles formed stomatocytes that detached from the membrane and floated away. Vesicles in the presence of 0.4% or 0.8% PEG experienced sufficient adhesion to allow deflation to reduced volumes as low as *ν <* 0.1 and rims with radii of curvature as low as *R*_*rim*_ *<* 15 nm (Fig. 7D and F). These values are estimates since the vesicle thickness and rim height were below the diffraction limit of the microscope. However, we could discern a slight upward bend of of the rims of the otherwise diffraction limited rims, suggesting that the GUVs remained intact even under such extreme degrees of deflation.

**Figure 7:**
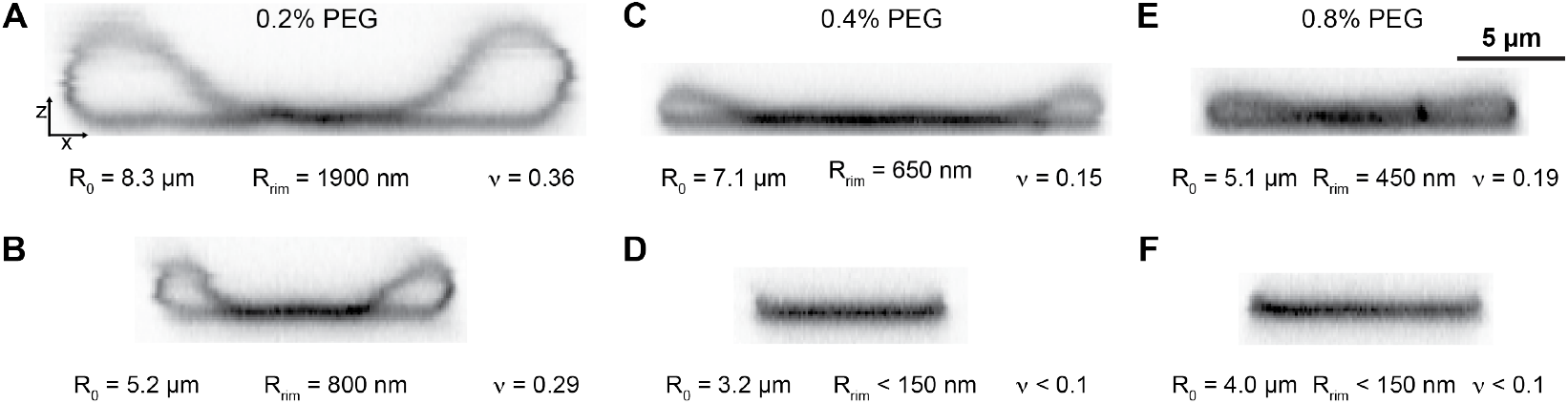
Bending-dominated shapes of highly deflated adhered vesicles. Color-inverted orthogonal cross-section of concave adhered vesicles (adhered discocytes) at adhesion strengths corresponding to PEG concentrations of **(A-B)** 0.2% (w/v), **(C-D)** 0.4% (w/v), and **(E-F)** 0.8% (w/v). Rows and columns show two different vesicle sizes and PEG concentrations, respectively.

## Conclusion

In summary, we have developed a simple and robust platform to modulate and characterize the shapes of adhered vesicles with reduced volumes as low as 0.1. We found that weak adhesion of the vesicles to a flat substrate was essential to retain stable shapes down to low reduced volumes, yielding curvature radii as small as *R*_*rim*_ = 150 nm at the rim and *R*_*top*_ = −200 nm at the top of the vesicle. Using shape analysis with the Canham-Helfrich model, we extracted key mechanical parameters, including adhesion strength, membrane tension, and vesicle pressure from the radial confocal slices of the GUVs. We found that the rate of shape flattening during deflation is determined by a normalized adhesion strength that combines vesicle size, adhesion energy, and bending rigidity. For flat, disc-like vesicles, we established a predictive relationship that allows the adhesion strength to be estimated solely based on the vesicle’s aspect ratio, size, and bending rigidity.

Future studies should explore how lipid composition influences the evolution of shape and mechanics of adhered vesicles accompanying deflation. While our results were obtained using POPC vesicle membranes, biological membranes feature diverse lipid mixtures with compositional asymmetries that can significantly alter curvature and mechanics. For instance, sphingomyelin content in the Golgi apparatus has been shown to affect cisternal morphology and function. ^65^ In addition, our shape analysis provides a novel approach to measure strength of membrane-membrane adhesion for a variety of systems including adhesion mediated by membrane-attached bio-molecular condensates, as found in tight junctions. ^66^ Extending our approach to GUVs obtained from octanol-assisted liposome assembly (OLA) or combined with synthetic membrane shapers could further enhance throughput and control over vesicle uniformity.^67,68^

Our findings have broader implications for understanding the mechanical constraints that govern the morphologies of membrane-bound organelles. In particular, they shed light on the adhesion strengths and membrane tensions required to achieve cisterna-like shapes *in vitro*. Membrane tension is a key parameter in vesicular trafficking, as it influences membrane fusion and the rate of cargo transport between compartments. Our modeling reveals how shape transitions are mechanistically coupled to tension and curvature, with cisterna-mimetic shapes appearing to exhibit ultra-low membrane tension, which may inhibit membrane fusion. Finally, the ability to dynamically control membrane curvature and compartment volume could be harnessed to develop novel nanomaterials with programmable mechanical and optical properties, or as microscale components in bioelectronic and energy transduction systems where surface-to-volume ratio is a critical design parameter.^69,70^

## Data Availability

The presented datasets are available from the corresponding author upon request.

## Acknowledgments

This work was supported by Ambizione Grant 202214 from the Swiss National Science Foundation to AAR. We thank Hendrik T. Spanke, Jan Steinkühler, Masayuki Imai, Peter Walde for insightful discussions. We thank Roland L. Knorr for helpful suggestions in the sample chamber design.

## Author Contributions Statement

AAR, GW, and RWS wrote the paper. AAR, RWS, GW, JAC, and ERD designed the experiments. GW performed the experiments. GW, RWS, JAC, AAR, ERD, and PB performed the data analysis.

