## Supporting Information for "Dynamic Shape Modulation of Deflated and Adhered Lipid Vesicles"

### Materials and Methods

#### GUV Preparation

Giant unilamellar vesicles (GUVs) were made by electroformation, following the protocol from Ref. 1, on indium tin oxide (ITO) coated glass plates (MSE Supplies, 10x10x1.1 mm with a sheet resistance of 3-5 Ohm/Sq (ITO layer thickness of 350 nm)).[1–5] We used a lipid composition of 99.9% (mol/mol) 1-palmitoyl-2-oleoyl-glycero-3- phosphocholine (POPC) (Avanti Polar Lipids, 850457C) and 0.1% (mol/mol) of 1,2-dioleoyl-sn-glycero3-phosphoethanolamine-N-(lissamine rhodamine B sulfonyl) (ammonium salt) (DOPE-Rh) (Avanti Polar Lipids, in chloroform, 810150C).

The electroformation chamber consisted of two ITO-coated glass plates separated by a 3 mm polydimethylsiloxane (PDMS) (Sigma Aldrich, Sylgard 184) spacer with cut-out wells of a 9 mm radius and small channels for filling. The lipid solutions were diluted in chloroform (Sigma Aldrich, 132950) to a final lipid concentration of 1 mM. We added dropwise 10 µL of the lipid solution to each well and let the chloroform evaporate. The residual chloroform was evaporated under vacuum overnight. The chamber was closed with the second ITO-coated glass plate, secured with clamps, and filled with GUV buffer. The GUV buffer consists of a filtered (TPP, 0.22 µm PES membrane) sucrose (Sigma Aldrich, S7903) solution at 25 mOsm/kg (GUV buffer), as determined by a freezing point osmometer (Osmomat 3000, Gonotec).

A function generator (Keysight, 33210A) was attached to the ITO plates through aluminium strips each side, resulting in an electric field perpendicular to the ITO plates. The AC voltage was increased in steps of 5 minutes to 5 V (0.83 V, 1.66 V, 2.5 V, 3.3 V, 4.1 V, 5 V) at a constant 10 Hz. The field was then kept for two hours at 5 V and 10 Hz and finally for 30 minutes at 5 V and 5 Hz. The vesicles were removed with glass pipettes and stored in glass tubes at 4° C, where they remained stable for weeks.

#### SUV and SLB Preparation

Supported lipid bilayers (SLB) were made by fusing small unilamellar vesicles (SUVs) onto glass coverslips (VWR, No. 1.5).[6] For the SUVs, 25 µL of 30 mM lipids with 99.9% (mol/mol) POPC (Avanti Polar Lipids, 850457C) and 0.1% (mol/mol) of 1,2-dioleoyl-sn-glycero-3-phosphoethanolamine-N-dibenzocyclooctyl (DOPE-DBCO) (Avanti Polar Lipids, 870129C) were dried to a film in a glass test tube under flushing argon gas. Residual chloroform was evaporated under vacuum overnight. The lipids were then resuspended in 1 mL SUV buffer by pipetting until the liquid was turbid. The SUV buffer consisted of 25 mM HEPES (Sigma Aldrich, H3375), 140 mM KCl (VWR, 26764.232), and Alexa Fluor 647 azide triethylammonium salt (Invitrogen, A10277) at a 1:1 molar ratio to the DOPE-DBCO lipids. The buffer was adjusted to pH 7.4 and filtered (TPP, 0.22 µm PES membrane). The solution was tip-sonicated (Branson, Sonifier 250) for 2-3 minutes at power 2 and output 20 % for 5-10 cycles with breaks of 5 minutes until the solution became transparent, taking care to avoid significant heating of the solution. Finally, the solution was centrifuged (Eppendorf, 5418 R) for 5 minutes at 16'000 g and the supernatant with SUVs was collected and stored at 4° C until used.

The glass coverslips were cleaned by immersing them in water and treating them in a bath sonicator (Fisherbrand, FB11201) for 10 minutes at 37 kHz at 100%. This was followed by immersion and sonication of the coverslips in ethanol. Next, the coverslips were rinsed with water, blow-dried with air, and surface-activated in a UV/Ozone cleaner (Bioforce Nanoscience, UV/Ozone Cleaner ProPlus) for 10 minutes. Immediately after UV/Ozone treatment, imaging spacers (SecureSeal, GRACE bio-labs, SS1X9) with 9 mm diameter and 120 µm depth were stuck onto the coverslips and the SLB was formed by adding 30-40 µL of SUV solution diluted 1:20 with SLB buffer to the spacer annulus. The SLB buffer consisted of 10 mM Tris pH 7.5 (Thermo Scientific, J63831), 150 mM NaCl (VWR, 27810.295), 2 mM $CaCl_2$ (Sigma Aldrich, 902179) in water and was filtered (TPP, 0.22 µm PES membrane) prior to use. This coverslip and spacer containing the SUV solution was incubated overnight and stored in a humidity box until used. Just before the experiment, residual SUV solution was removed by washing the SLB with 3 mL of outside buffer (10 mM NaCl (VWR, 27810.295) and 5 mM glucose (Sigma Aldrich, G7528)) and then with 3 mL outside buffer including 100 kDa polyethylene glycol (PEG) (Sigma Aldrich, 181986) at the final concentration of 0.17% (w/v), 0.34% (w/v) or 0.84% (w/v). The PEG concentration after filtration (TPP, 0.22 µm PES membrane) was determined by $^1$H NMR spectroscopy. Deuterated water ($D_2O$) (Apollo Scientific, DE50B) was used as the solvent and dimethylformamide (DMF) (Fisher Scientific, D/3840/17) was used as a concentration standard as previously described.[7] The PEG concentration was determined based on the ratio of the integrated peaks corresponding to DMF (at known concentration) and the PEG.

#### Diffusion Chamber for Deflation at Constant Adhesion Strength

The SLB covered with PEG-containing outside buffer was placed on the microscope and 1 or 10 µL GUV solution were slowly added, subsequently letting the GUVs sediment for 5 to 20 minutes. Meanwhile, a dialysis membrane (3.5 kDa Mini Dialysis Device 2 mL, SlideA-Lyzer) was washed with water and equilibrated with outside buffer. The empty dialysis membrane was then carefully placed onto the spacer and 2 mL of outside buffer were added inside the dialysis button. The dialysis membrane was permeable to water, glucose, and NaCl, but not to PEG 100 kDa.

The system was left to equilibrate for 1 hour. Meanwhile, 10-15 microscope stage positions were selected where vesicles were visible, and confocal z-stacks were acquired (z-step size 0.27 µm) using a spinning disc confocal microscope (Nikon Eclipse Ti2 base with Yokogawa CU-W1 with XYZ automated stage with piezo Z-axis PZ-2300 from Applied Scientific Instrumentation) on a 60x oil objective (Nikon, NA 1.4). The vesicle and SLB membranes were imaged with the 560 nm and 640 nm excitation laser lines, respectively. After imaging of each deflation step, 50 µL of the solution inside the dialysis cup was extracted and its osmolarity was measured using a freezing point osmometer (Osmomat 3000, Gonotec). To achieve a homogeneous concentration in the dialysis membrane, around 800 µL solution was removed, externally mixed with 50 uL deflation solution, added back, and mixed by pipetting up and down. At each deflation step, the composition and osmolarity of our system equlibrated within an hour via diffusion across the dialysis membrane. To obtain the desired resolution of different shapes through deflation, this process was repeated 4 times, while we increased the amount of added deflation buffer from 50 µL to 100 µL, 100 µL, and 200 µL. The total volume in the dialysis cup was kept constant at 2 mL.

#### Image Processing

The SLB and GUV adhesion zone are in the xy-plane. The stacks are sliced in xz to obtain an orthogonal sliced vesicle cross-section. We correct for a z-aberration from the use of an oil-immersion objective by a factor of 0.878, resulting in a z-step of 0.237. First, we identify single GUVs over all time points by location. Then the vesicle is vertically sliced through the center into 3 orthogonal xz cross-sections. With the 3 slices, we estimate the deviation from perfect radial symmetry and create error bars.

#### Shape Fitting with Canham-Helfrich Model

The shapes and accompanying mechanical parameters were found by solving the Canham-Helfrich model with adhesion following previously described procedures.[8,9] Only axisymmetric shapes were considered. These shapes could therefore be described as surfaces of revolution around the z-axis of a contour parametrized by the arc-length $s$. The contour was determined by the coordinates perpendicular and parallel to the axis of symmetry, ($X(s)$, $Z(s)$) with the accompanying geometric relations (Fig. S9),

$$\frac{\mathrm{d}X}{\mathrm{d}s} = \cos\psi \tag{S1}$$

$$\frac{\mathrm{d}Z}{\mathrm{d}s} = -\sin\psi \tag{S2}$$

$$C_1 = \frac{\mathrm{d}\psi}{\mathrm{d}s} \tag{S3}$$

$$C_2 = \frac{\sin\psi}{X}, \tag{S4}$$

where $\psi$ is the tilt angle of the surface with respect to the x-axis, and $C_1$ and $C_2$ are the principal curvatures. The vesicle shape minimizes the free energy,

$$F = \underbrace{\frac{\kappa}{2}\oint (C_1 + C_2)^2 \,\mathrm{d}A}_{E_{\text{bending}}} + \underbrace{\sigma A}_{E_{\text{tension}}} + \underbrace{\Delta P \cdot V}_{E_{\text{pressure}}} - \underbrace{\omega A_{adh}}_{E_{\text{adhesion}}}, \tag{S5}$$

where $\kappa$ is the bending rigidity, $\sigma$ is the membrane tension, $A$ is the membrane surface area, $\Delta P = P_{in} - P_{out}$ is the vesicle pressure, $V$ is the volume of the vesicle, $\omega$ is the adhesion strength, and $A_{adh}$ is the adhered area. Using the parametrization defined by Eqs. S1-S4 yields the energy functional

$$F\left[X(s), \psi(s), s\right] = 2\pi \int_0^{s_1} L\left(X, \dot{X}, \psi, \dot{\psi}, \gamma\right) \mathrm{d}s, \tag{S6}$$

where $L\left(X, \dot{X}, \psi, \dot{\psi}, \gamma\right)$ is the Lagrange function

$$L\left(X, \dot{X}, \psi, \dot{\psi}, \gamma\right) = \frac{\kappa}{2}\left(\frac{\mathrm{d}\psi}{\mathrm{d}s} + \frac{\sin\psi}{X}\right)^2 X + \sigma X \,\mathrm{d}s + \frac{PX^2}{2}\sin\psi + \gamma \cdot \left(\frac{\mathrm{d}X}{\mathrm{d}s} - \cos\psi\right) \tag{S7}$$

and $s_1$ is the arc length at which the shape meets the surface with

$$\psi(s_1) = \pi \tag{S8}$$

as shown in Fig. S9, and $\gamma = \gamma(s)$ is a Lagrange parameter function to enforce the geometric constraint, $\dot{X} = \cos\psi$. This parametrization divides the shape into a free portion (Fig. 5 A-C *green*), which is derived from numerical integration, and a flat adhered portion (*blue*), which is defined implicitly as a disc with radius $X(s_1)$. It is useful to adopt the non-dimensionalized variables, $\overline{X} = X \cdot U_0$, $\overline{Z} = Z \cdot U_0$, and $\overline{s} = s \cdot U_0$, where $U = \dot{\psi}$ is the principal curvature curvature along the contour, $C_1$, and $U_0 = |U(s=0)| = 1/|R_{top}|$, with $R_{top}$ the signed radius of curvature at the top of the vesicle. Minimization of the energy functional, $F$ (Eq. S6), yields the non-dimensionalized Euler-Lagrange equations,

$$\frac{\mathrm{d}\psi}{\mathrm{d}\overline{s}} = \overline{U} \tag{S9}$$

$$\frac{\mathrm{d}\overline{U}}{\mathrm{d}\overline{s}} = -\frac{\overline{U}}{\overline{X}}\cos(\psi) + \frac{\cos(\psi)\sin(\psi)}{\overline{X}^2} + \frac{\overline{\gamma}}{\overline{X}}\sin(\psi) + \frac{\Delta\overline{P}\overline{X}}{2}\cos(\psi) \tag{S10}$$

$$\frac{d\overline{\gamma}}{d\overline{s}} = \frac{\overline{U}^2}{2} - \frac{\sin^2(\psi)}{2\overline{X}^2} + \Delta\overline{P} \cdot \overline{X}\sin(\psi) + \overline{\sigma}, \tag{S11}$$

and

$$\frac{d\overline{X}}{d\overline{s}} = \cos(\psi), \tag{S12}$$

We then numerically solve these equations using initial conditions

$$\overline{X}(\overline{s} = 0) = 0 \tag{S13}$$

$$\psi(\overline{s} = 0) = 0 \tag{S14}$$

$$\overline{\gamma}(\overline{s} = 0) = 0 \tag{S15}$$

and

$$\overline{U}_0 = \overline{U}(\overline{s} = 0) = \pm 1\,, \tag{S16}$$

whereby the last equation defines the direction of the curvature, with $\overline{U}_0 = 1$ for convex and $\overline{U}_0 = -1$ for concave vesicle tops, respectively. The three parameters that control the shape of the vesicle are the normalized tension $\overline{\sigma} = \sigma \cdot R_{top}^2/\kappa$, the normalized pressure $\Delta\overline{P} = \Delta P \cdot R_{top}^3/\kappa$, and the initial condition of convex or concave $\overline{U}_0$ (Eq. S16).

The equations were solved using the ode45 function in MATLAB and by integrating from the top of the vesicle until $s_1$ defined by Eq. S8. The adhesion strength, was determined from the curvature at the contact line (Fig. 2A),[9]

$$\frac{1}{\overline{R}_c} = \dot{\psi}\,(\overline{s}_1) = \sqrt{2\overline{\omega}}\,. \tag{S17}$$

The dimensionful mechanical parameters were obtained by using $\kappa = 33\,\mathrm{k_B T}$ for POPC membranes and the best-fit $U_0$.[1]

The calculated curves were overlaid on the experimentally observed radial slices using a custom MATLAB GUI. First, the axes of symmetry of the experimentally observed vesicle and the numerically calculated shape were aligned. Next, the maximum radius, $R_{max}$ (Fig. 2A), of the numerically calculated shape was adjusted so that maximum widths of the calculated shape and the observed shape coincided.

Vesicles approximating spherical caps (Fig. 5A) were fitted with convex top curvature, $\overline{U}_0 = 1$, and overpressure, $\Delta\overline{P} > 0$. A ratio $\Delta\overline{P}/\overline{\sigma} = 2 - \varepsilon$, with $\varepsilon << 1$, yielded spherical cap-like shapes, whereby greater absolute values of $\overline{\sigma}$ and $\Delta\overline{P}$ decreased the reduced volume and contact angle of the shape (see S12). Decreasing the ratio $\Delta\overline{P}/\overline{\sigma}$ increasingly flattened the shapes, culminating in perfectly flat shapes for $\Delta\overline{P} = 0$. Accordingly, flat disc-like shapes were fitted with a convex top curvature with $\Delta\overline{P} \approx 0$ (Fig. 5B). However, concave top curvature and $\Delta\overline{P} \ll 0$ also yielded flat shapes (Fig. S13). In either case, the aspect ratio and height of the disc is controlled by $\overline{\sigma}$. Lower heights (smaller $R_{rim}$) were achieved by increasing $\overline{\sigma}$.

Concave adhered discs were fitted with concave top curvature, $\overline{U}_0 = -1$, and underpressure, $\Delta\overline{P} < 0$. Decreasing $\Delta P < 0$ reduced the curvature at the top of the vesicle, $U_0$, culminating in $U_0$ for $\Delta P \ll 0$. Similar to the disc-like shapes, $R_{rim}$ could be matched to observations by adjusting $\overline{\sigma}$. Importantly, this fitting method works on adhered shapes that show impact from bending on the shape. It fails when fitting spherical caps in the strong adhesion limit, where the bending does not impact the shape, which is needed for correct scaling of the energies. Furthermore, for $\Delta P < 0$ and $\Delta\overline{P}/\overline{\sigma} = -(2 + \varepsilon)$, the model yields unphysical predictions where the top of the membrane penetrates the adhered portion of the vesicle.

To obtain numerical predictions for $-0.25 < C_{top}R_0 < 0.25$ (Figs. 5F and 6C-F), the shape equations were solved using the shooting method from the top of the vesicle and by setting as initial condition $U_0 = C_{top} = 0$. The unknown value of $\sigma R_0^2/\kappa$ was used to satisfy the total area constraint, $A/(4\pi R_0^2) = 1$, for some fixed $\Delta P R_0^3/\kappa$. This gives a shape for $C_{top} = 0$ with a certain reduced volume. This was then repeated over a range of values of $\Delta P R_0^3/\kappa$, yielding a family of shapes with $C_{top} = 0$ and possessing various reduced volumes.

The same process was used to obtain shapes with $C_{top}R_0 = -0.25$ and $C_{top}R_0 = 0.25$.

#### Quantification of Shape Geometries

The vesicle area was calculated by integrating $X(s)$ from the best-fit shape,

$$A = \int_0^{s_1} 2\pi X \,\mathrm{d}s \tag{S18}$$

and the volume was determined using

$$V = \int_0^{s_1} 2\pi X^2 \sin(\psi)\,\mathrm{d}s\,. \tag{S19}$$

The purely geometric quantities (Figs. 3 and 4), including vesicle area, volume, reduced volume, height, maximum radius, and vesicle top radius of curvature were obtained for all vesicles (including spherical caps where shape analysis was not feasible) using the manual contour tracking method previously reported by Steinkühler et. al.[10]

The reduced volume $\nu$ was calculated using

$$\nu = \frac{V}{V_0} = \frac{V}{\frac{4\pi}{3}\cdot R_0^3} = 6\sqrt{\pi}\frac{V}{A^{3/2}}, \tag{S20}$$

where $V$ is the volume of the deflated object, $V_0$ is the volume of a sphere with the same surface area as the object. Therefore, lower reduced volumes indicate a higher degree of deflation or respectively a larger surface-to-volume ratio. The reduced volume of a sphere is $\nu_0 = 1$.

#### Bendocapillary Length and Rim Height

The relationship $R_{rim} \approx \lambda_\kappa$ with $\lambda_\kappa \equiv \sqrt{\kappa/\sigma}$ the bendocapillary length was derived following Forêt et al.[11] Briefly, the shape of the highly curved rim of an axisymmetric vesicle is well-described by the shape of a 2-dimensional contour that extends infinitely into the plane of the contour. This approximation holds as long as the equatorial curvature along the widest part of the vesicle, $C_2 = \sin\psi/X$, which is at most of order $O(1/R_0)$, is negligible relative to the rim curvature $C_1 = \dot{\psi}$, which is of order $O(1/\lambda_\kappa)$. The rim of a flat disc-like vesicle then forms syntractrix given by[11]

$$x(z) = \lambda_\kappa \ln\left(\frac{2\lambda_\kappa + \sqrt{4\lambda_\kappa^2 - z^2}}{z}\right) - \sqrt{4\lambda_\kappa^2 - z^2}, \tag{S21}$$

which is defined for $0 < z \leq 2\lambda_\kappa$. In particular, this $z$-range implies that $h = 2\lambda_\kappa$ and therefore that $R_{rim} = h/2 = \lambda_\kappa$.

#### Supplementary Figures

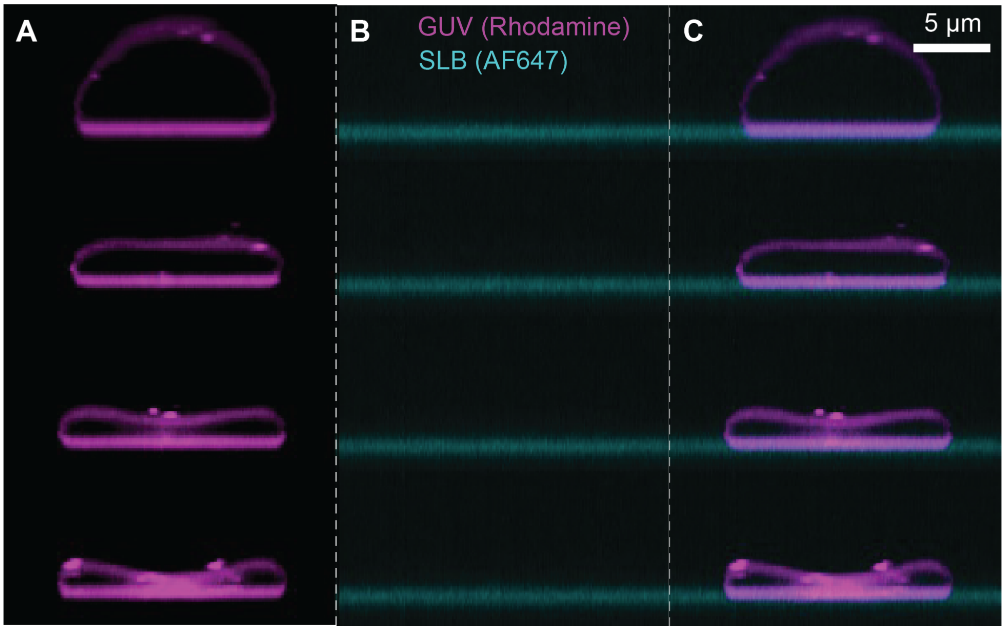


Figure S1: *Two-color confocal imaging of fluorescently labelled lipids in the GUV and supported lipid bilayer (SLB) membranes.* Orthogonal xz cross-sections show a single GUV (*magenta*) on a SLB (*cyan*) undergoing deflation. **(A)** GUVs containing rhodamine-DOPE lipids. **(B)** SLB containing DBCO-DOPE lipids labelled with AlexaFluor647-azide using copper-free click chemistry. **(C)** Overlay of GUV and SLB channels. The SLB and GUV membranes do not exchange lipids over the course of adhesion and deflation, indicating absence of hemi-fusion or full fusion between the two membranes.

Table S1: *Parameters for the best-fit axisymmetric vesicle shapes from Fig. 5.* Columns correspond to vesicles shown in Fig. 5 (A), (B), and (C). The top 4 rows list the dimensionless parameters for numerical integration of the shape equations (Eqs. S9 - S12), where $\Delta\overline{P} = \Delta P \cdot R_{top}^3/\kappa$, $\overline{\sigma} = \sigma \cdot R_{top}^2/\kappa$, and $R_{max}$ is the maximum disc radius. Subsequent rows list the corresponding dimensionful quantities assuming $\kappa = 33\, k_B T$.

| | **A** | **B** | **C** |
|---|---|---|---|
| $\Delta\overline{P}$ | 399.94 | 0 | -1000 |
| $\overline{\sigma}$ | 200 | $10^6$ | 484 |
| $R_{max}$ ($\mu$m) | 11.5 | 11 | 11 |
| Curvature direction | convex | convex | concave |
| $R_{top}$ ($\mu m$) | 16 | 1358 | 30 |
| $\omega$ ($k_BT/\mu$m$^3$) | 48 | 45 | 47 |
| $\sigma$ ($k_BT/\mu$m$^2$) | 27 | 18 | 18 |
| $\Delta P$ ($k_BT/\mu$m$^3$) | 3.5 | 0 | -1.2 |
| $A$ ($\mu$m$^2$) | 1018 | 852 | 838 |
| $A_{adh}$ ($\mu$m$^2$) | 360 | 338 | 340 |
| V ($\mu$m$^3$) | 1900 | 926 | 625 |
| $\nu$ | 0.64 | 0.40 | 0.27 |
| $A_{adh}/A$ | 0.35 | 0.40 | 0.40 |

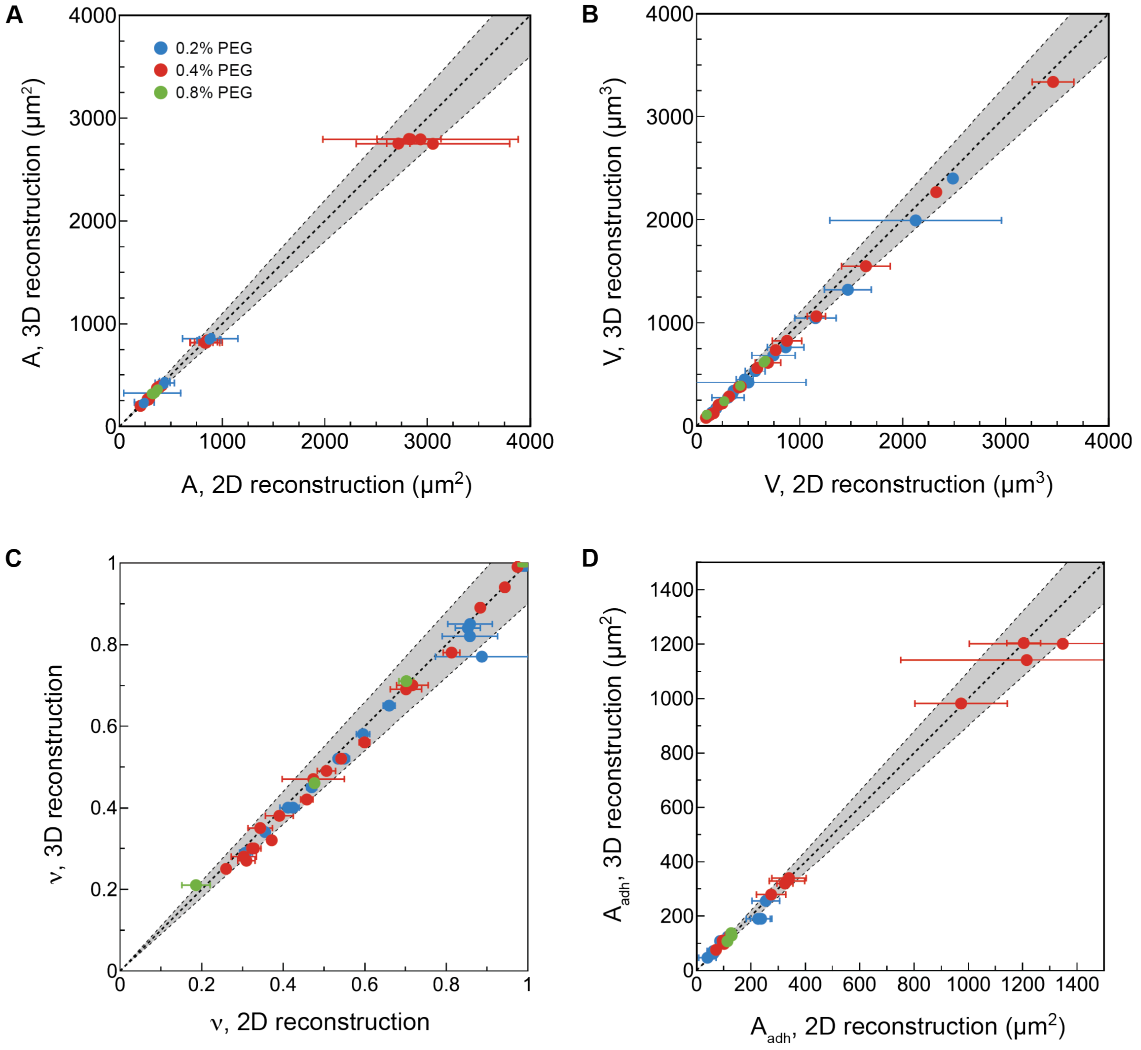


Figure S2: *Comparison between geometric observables obtained from three-dimensional reconstruction and a simplified approach, assuming axiosymmetry and inference from a two-dimensional contour within a radial slice of the vesicle.* The two methods yield the same results within 10% accuracy (gray shaded area) for **(A)** the membrane surface area $A$, **(B)** the enclosed volume $V$, **(C)** the reduced volume $\nu$, and **(D)** the adhered area $A_{adh}$. The error bars represent the standard deviation among measurements from three radial slices of a vesicle. For the 3D reconstruction, a custom python script was used. Colors indicate the concentration of PEG 100 kDa in the outside buffer: 0.2% (w/v) (*blue*), 0.4% (w/v) (*red*), 0.8% (w/v) (*green*).

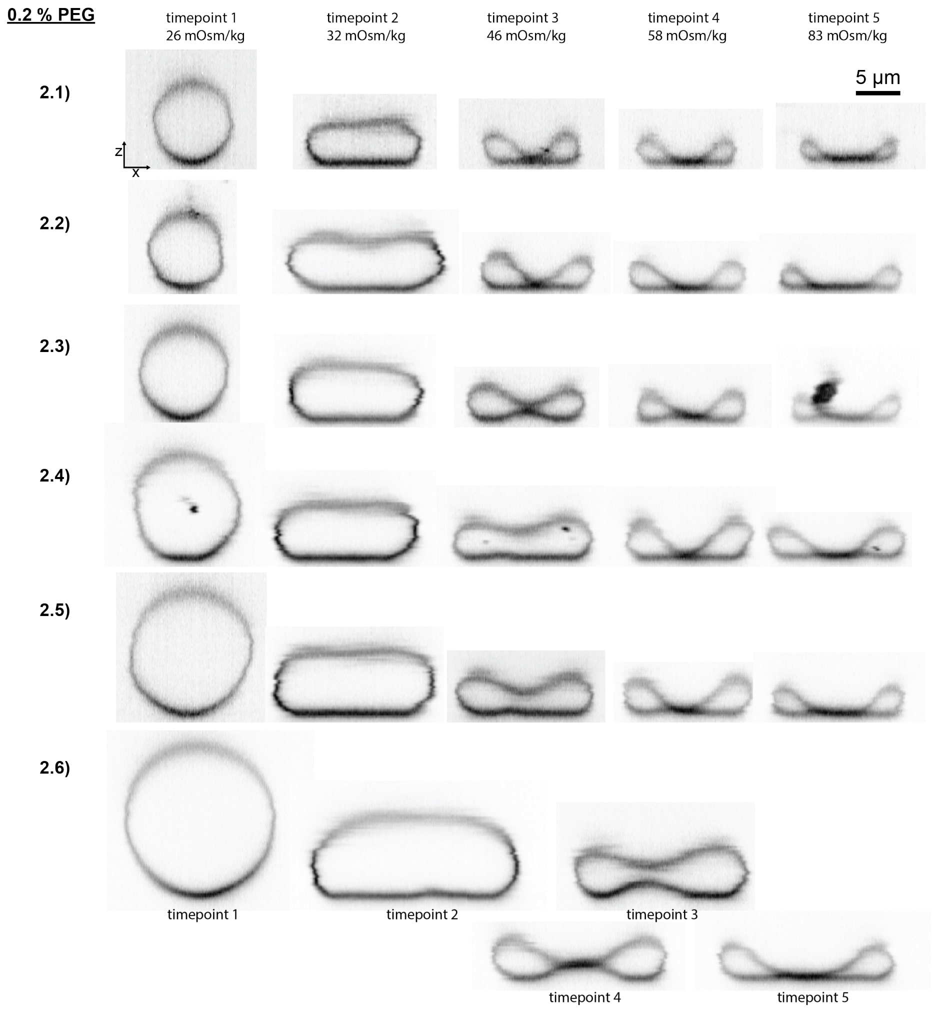


Figure S3: *Radial slices from confocal z-stacks of vesicles at 0.2 % (w/v) PEG sorted by increasing size.* Each row corresponds to one vesicle imaged over the course of the deflation sequence. Scale bar: 5 um.

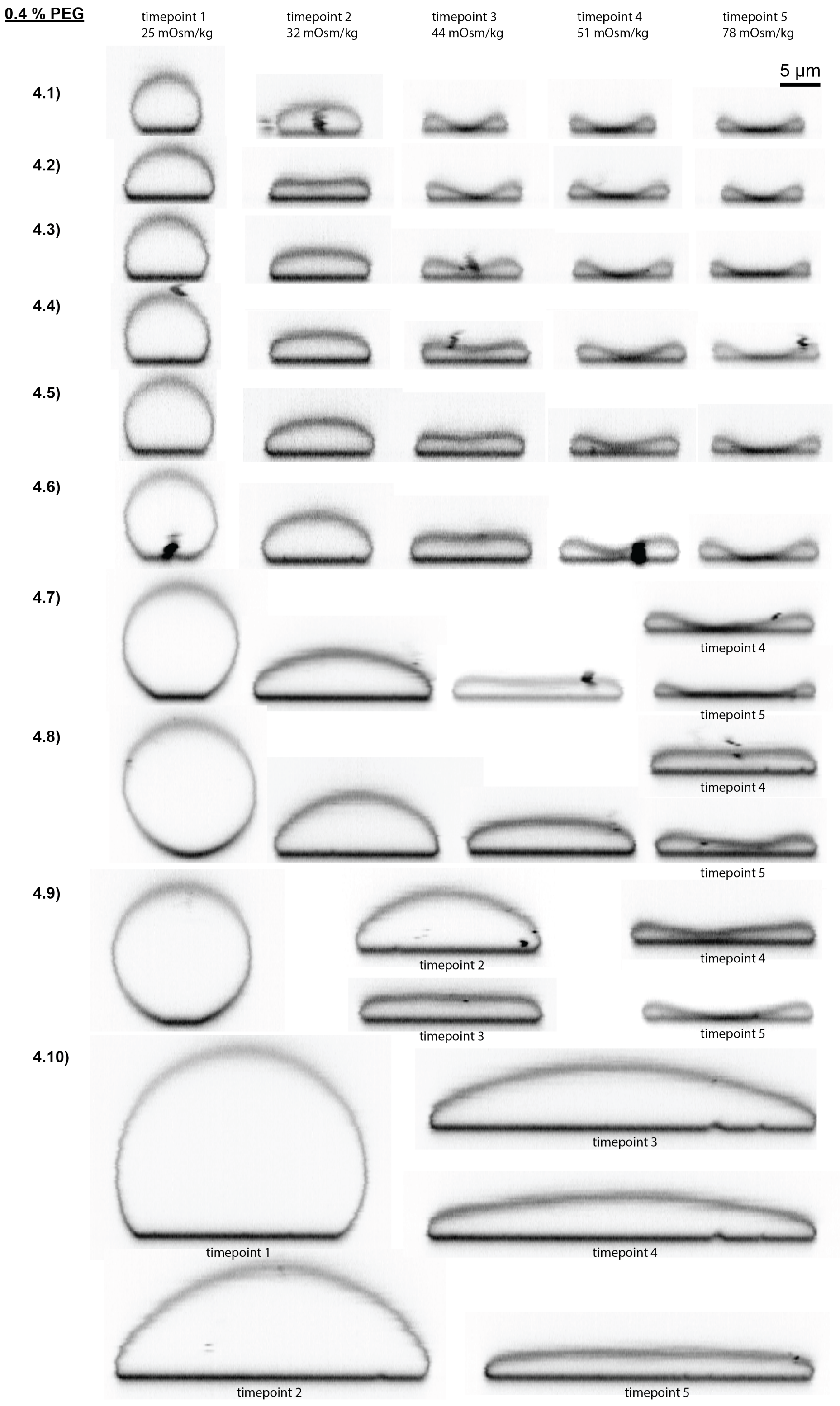


Figure S4: *Radial slices from confocal z-stacks of vesicles at 0.4 % (w/v) PEG sorted by increasing size.* Each row corresponds to one vesicle imaged over the course of the deflation sequence. Scale bar: 5 um.

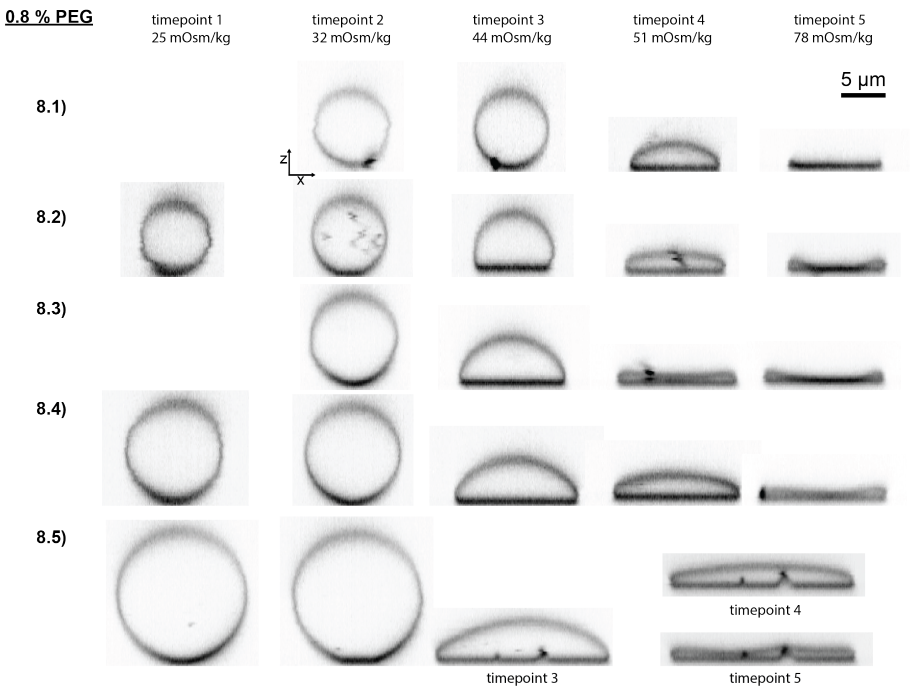


Figure S5: *Radial slices from confocal z-stacks of vesicles at 0.8 % (w/v) PEG sorted by increasing size.* Each row corresponds to one vesicle imaged over the course of the deflation sequence. Scale bar: 5 um.

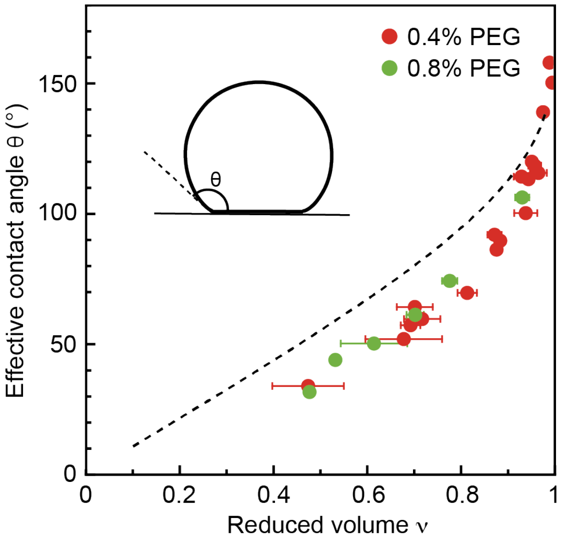


Figure S6: *Effective contact angle of spherical cap-like shapes as a function of reduced volume.* The effective contact angle $\theta$ was determined using the ImageJ Angle tool. Black dashed curve denotes the effective contact angle predicted for perfect spherical caps in the limit of infinitely strong adhesion (Eq. 2). The colors indicate the concentration of PEG 100 kDa in the outside buffer: 0.4% (w/v) (*red*) 0.8% (w/v) (*green*). Error bars indicate standard deviations among measurements obtained from 3 different confocal slices (see Methods and Materials for details).

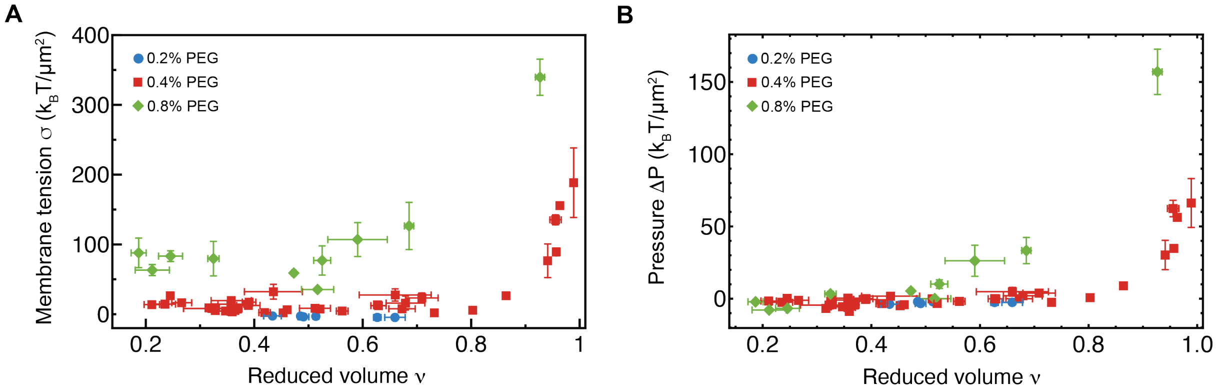


Figure S7: *Membrane tension and pressure from shape analysis* **(A)** Membrane tension, $\sigma$, as a function of reduced volume. **(B)** Pressure difference,m $\Delta P$, as a function of reduced volume.

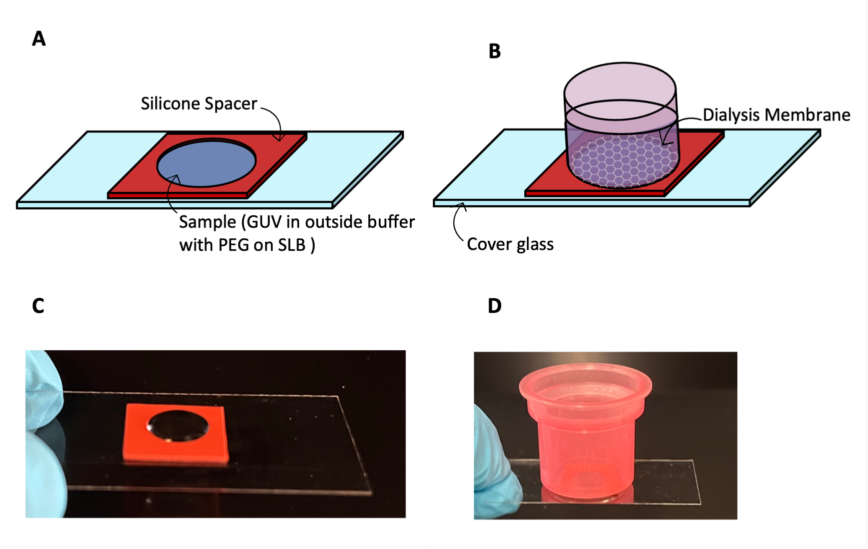


Figure S8: *Diffusion chamber for observation of individual GUVs during successive steps of osmotic deflation* **(A)** Schematic showing the imaging chamber. The supported lipid bilayer (SLB) was generated on the cover glass within the annulus of the silicone spacer. The annulus was then filled with a solution of GUVs in outside buffer containing a fixed concentration of PEG. **(B)** After the GUVs were allowed to sediment, a disposable dialysis cup was filled with deflation buffer and carefully placed on top of the spacer, bringing the dialysis membrane in direct contact with the GUV solution. **(C)** Photograph of the silicone spacer shown in schematic (A). **(D)** Photograph of the diffusion chamber with the dialysis cup placed on top of the silicone spacer, as shown in schematic (B).

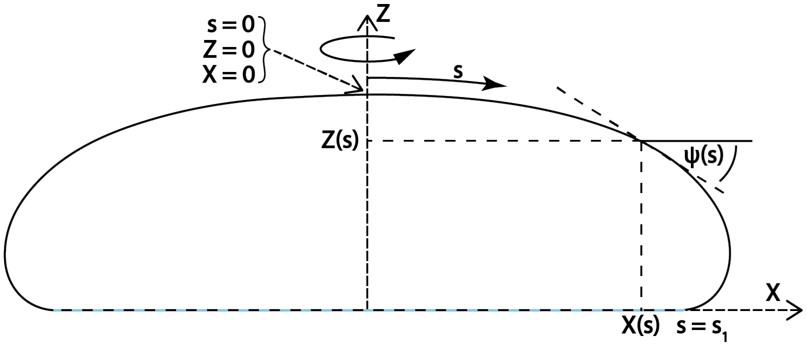


Figure S9: *Parametrization of the adhered vesicle shape used in the modeling.* The shape is an orthogonal xz cross-section, with rotational symmetric around the z axis. The shape is defined by the arclength S and the its slope $\psi$. The adhered membrane starts at an arclength $S = S_1$ and is marked by a blue-black dashed line.

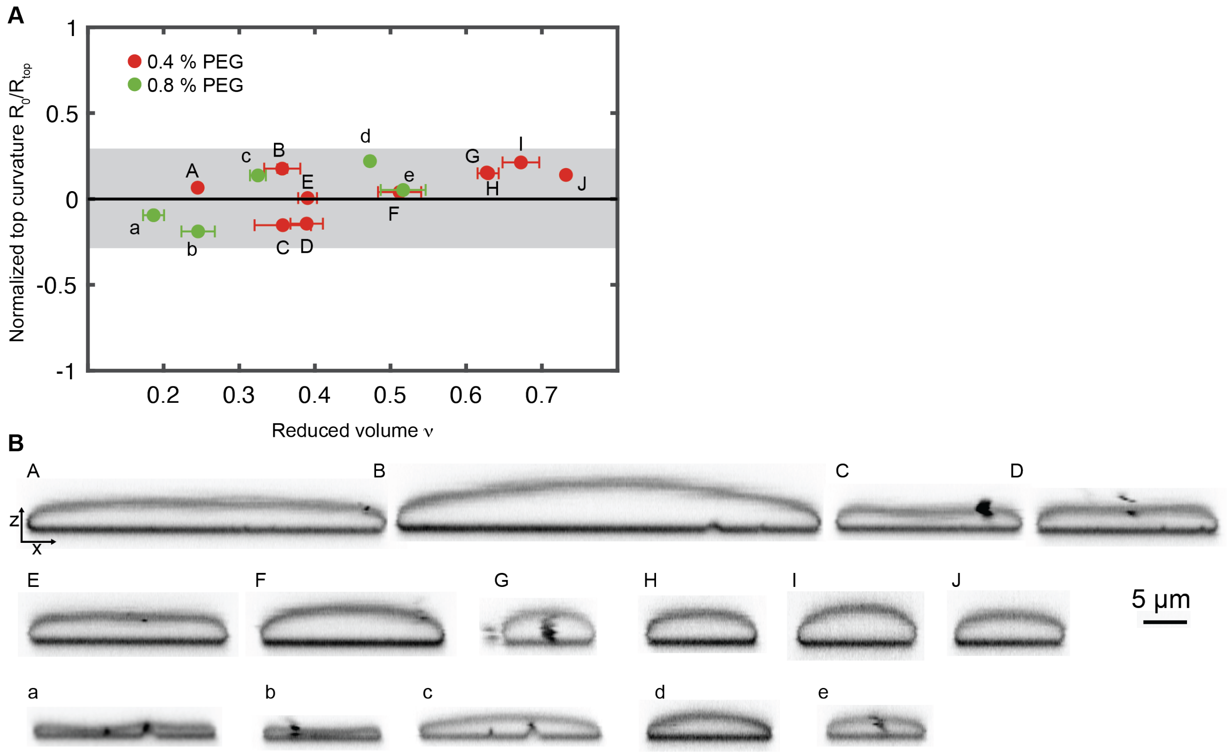


Figure S10: *Overview of flat disc-like vesicles shown in Fig.* 6 **(A)** Normalized to top curvatures and reduced volumes of the flat disc-like vesicles along with their labels at 0.4% (w/v) PEG (*capital letters*) and 0.8%v (w/v) PEG (*small letters*). **(B)** radial slices of the flat disc-like vesicles with labels as defined in (A).

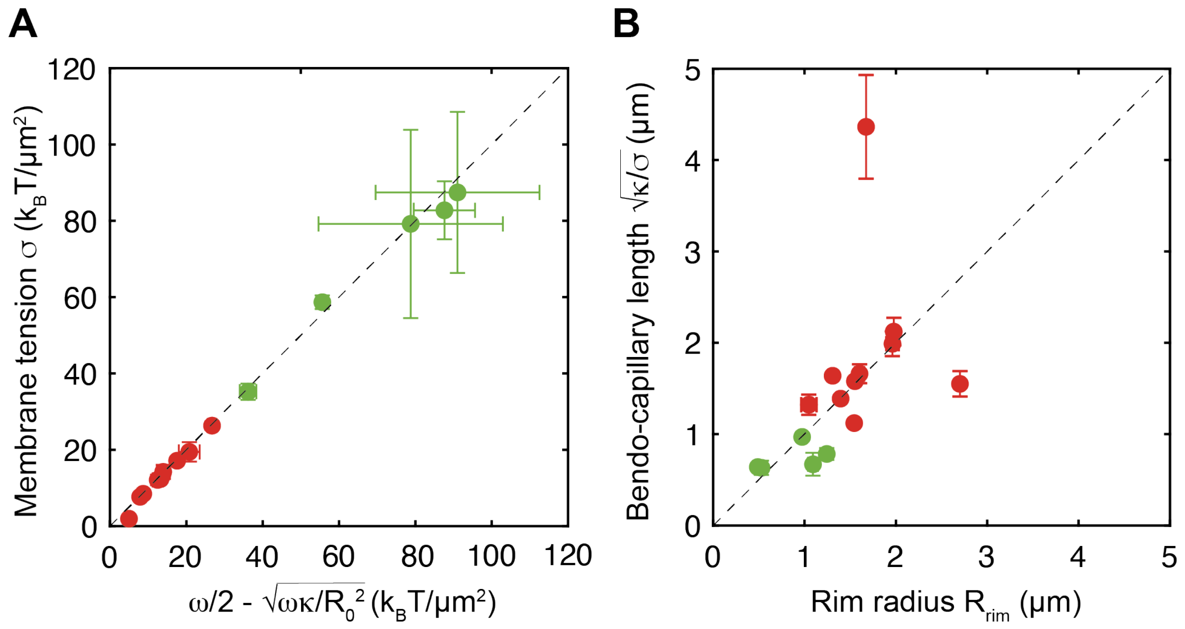


Figure S11: *Variation of Fig. 6 D and E with dimensions.* **(A)** The dimensional modified Young's law sets the tension, addition to Fig. 6 D. **(B)** The vesicle height is set by the bendo-capillary length, Fig. 6 E.

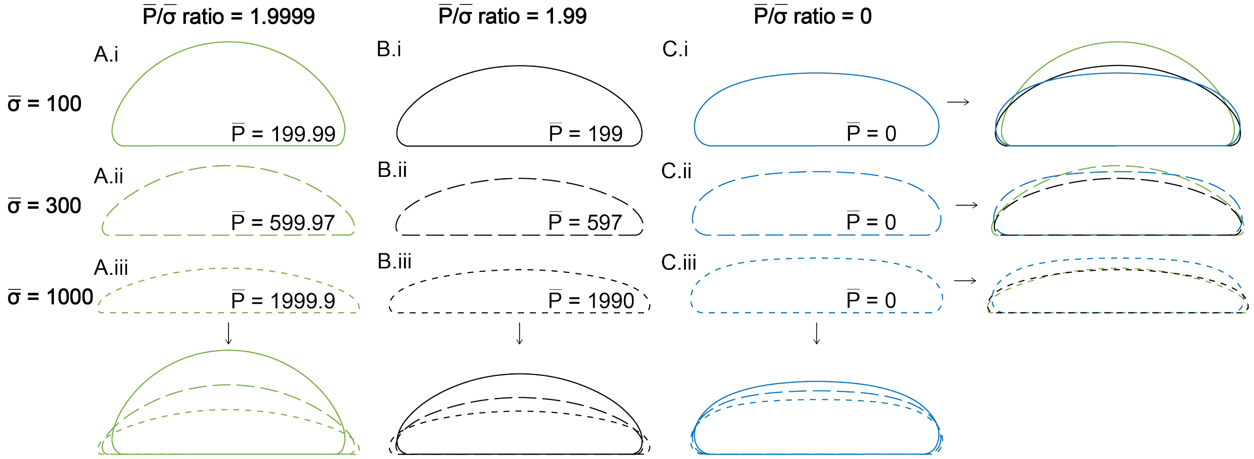


Figure S12: *Convex to disc-like shapes calculated by numerically integrating the shape equations of the Canham-Helfrich model.* Each of the three rows *(i)*-*(iii)* corresponds to a non-dimensionalized tension $\overline{\sigma} = \sigma \cdot R_{top}^2/\kappa$, with *(i)* $\overline{\sigma} = 100$, *(ii)* $\overline{\sigma} = 300$, and *(iii)* $\overline{\sigma} = 1000$. Each of the three columns *(A)*-*(C)* corresponds to a particular pressure-to-tension ratio with *(A)* $\overline{P}/\overline{\sigma} = 1.9999$ *(B)* $\overline{P}/\overline{\sigma} = 1.99$, and *C)* $\overline{P}/\overline{\sigma} = 0$. The lowest row and the right-most column show overlaid shapes. Comparing the shapes within a column, reveals that increasing $\overline{\sigma}$ decreases the aspect ratio and and the reduced volume. Comparing the shapes within a row, a decrease in $\Delta\overline{P}/\overline{\sigma}$ decreases increases $R_{top}$ and decreases the curvature at the rim. All vesicles are rescaled to a total surface area of $A = 1000\,\mu\text{m}^2$. Extracted parameters are shown in Table S2.

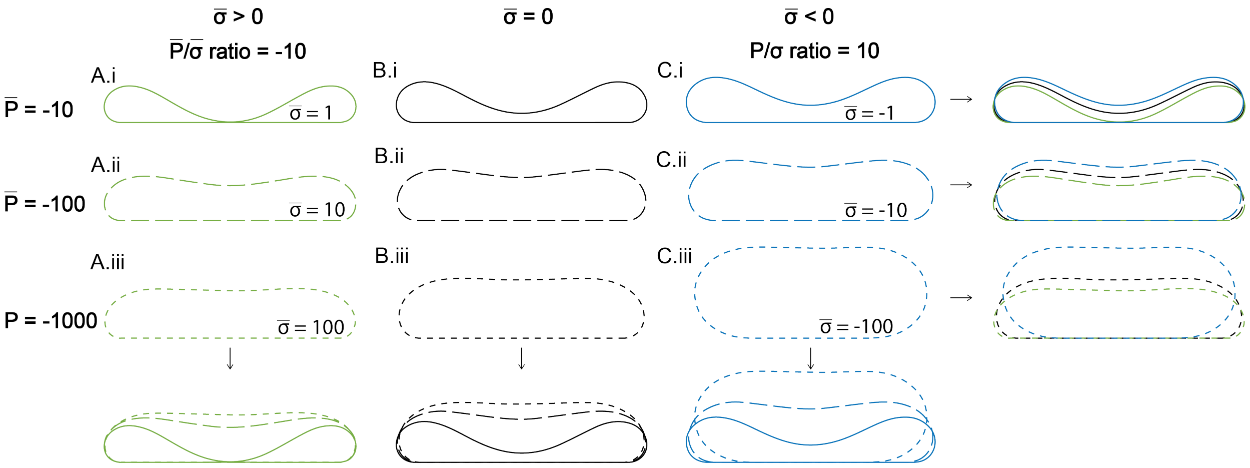


Figure S13: *Concave disc-like vesicle shapes calculated by numerically integrating the shape equations of the Canham-Helfrich model.* Each of the three rows *(i)-(iii)* corresponds to a non-dimensionalized pressure difference $\Delta\overline{P} = \Delta P \cdot R_{top}^3/\kappa$, with *(i)* $\Delta\overline{P} = 10$, *(ii)* $\Delta\overline{P} = 100$, and *(iii)* $\Delta\overline{P} = 1000$. Each of the three columns *(a)-(c)* corresponds to a particular pressure-to-tension ratio with *(a)* $\overline{P}/\overline{\sigma} = 10$ *(b)* $\overline{P}/\overline{\sigma} = 0$, and *c)* $\overline{P}/\overline{\sigma} = -10$. The lowest row and the right-most column show overlaid shapes. Comparing the shapes within a column, reveals that increasing $\Delta\overline{P}$ decreases $R_{top}$ and decreases the curvature at the rim. Comparing the shapes within a row, a decrease in $\Delta\overline{P}/\overline{\sigma}$ decreases the aspect ratio and increases the reduced volume. All vesicles are rescaled to a total surface area of $A = 1000\,\mu\text{m}^2$. Extracted parameters are shown in Table S3.

Table S2: *Parameters for the calculated shapes in Fig. S12 solving the Canham-Helfrich model.* Columns correspond to vesicles shown in Fig. S12. The top 4 rows list the dimensionless parameters for numerical integration of the shape equations (Eqs. S9 - S12), $\Delta\overline{P} = \Delta P \cdot R_{top}^3/\kappa$, $\overline{\sigma} = \sigma \cdot R_{top}^2/\kappa$, $R_{max}$ is the maximum disc radius, and curvature direction is denoted by "-" for concave and "+" for convex shapes. Subsequent rows list the corresponding dimensionful quantities assuming $\kappa = 33\,k_B T$.

| | A.i | A.ii | A.iii | B.i | B.ii | B.iii | C.i | C.ii | C.iii |
|---|---|---|---|---|---|---|---|---|---|
| $\Delta\overline{P}$ | 199.99 | 599.97 | 1999.9 | 199 | 597 | 1990 | 0 | 0 | 0 |
| $\overline{\sigma}$ | 100 | 300 | 1000 | 100 | 300 | 1000 | 100 | 300 | 1000 |
| $R_{max}$ ($\mu$m) | 10.64 | 11.52 | 11.95 | 11.19 | 11.29 | 11.88 | 11.17 | 11.35 | 11.50 |
| Curvature direction | + | + | + | + | + | + | + | + | + |
| $R_{top}$ ($\mu m$) | 11.51 | 16.29 | 27.42 | 16.04 | 23.34 | 39.48 | 35.59 | 52.47 | 82.49 |
| $\omega$ ($k_BT/\mu$m$^3$) | 20.50 | 40.91 | 55.31 | 14.97 | 23.75 | 29.22 | 5.51 | 7.00 | 8.88 |
| $\sigma$ ($k_BT/\mu$m$^2$) | 14.70 | 22.02 | 25.89 | 7.56 | 10.72 | 12.49 | 1.54 | 2.12 | 2.86 |
| $\Delta P$ ($k_BT/\mu$m$^3$) | 2.56 | 2.70 | 1.89 | 0.94 | 0.91 | 0.63 | 0 | 0 | 0 |
| $A$ ($\mu$m$^2$) | 1000 | 1000 | 1000 | 1000 | 1000 | 1000 | 1000 | 1000 | 1000 |
| $A_{adh}$ ($\mu$m$^2$) | 290 | 373 | 412 | 323 | 347 | 394 | 288 | 312 | 332 |
| V ($\mu$m$^3$) | 2323 | 1771 | 1320 | 2036 | 1518 | 1370 | 2062 | 1902 | 1746 |
| $\nu$ | 0.78 | 0.60 | 0.44 | 0.68 | 0.56 | 0.46 | 0.69 | 0.64 | 0.59 |
| $A_{adh}/A$ | 0.29 | 0.37 | 0.41 | 0.32 | 0.37 | 0.39 | 0.29 | 0.31 | 0.33 |

Table S3: *Parameters for the calculated shapes in Fig. S13 solving the Canham-Helfrich model.* Columns correspond to vesicles shown in Fig. S13. The top 4 rows list the dimensionless parameters for numerical integration of the shape equations (Eqs. S9 - S12), $\Delta\overline{P} = \Delta P \cdot R_{top}^3/\kappa$, $\overline{\sigma} = \sigma \cdot R_{top}^2/\kappa$, $R_{max}$ is the maximum disc radius, and curvature direction is denoted by "-" for concave and "+" for convex shapes. Subsequent rows list the corresponding dimensionful quantities assuming $\kappa = 33\, k_B T$.

| | a.i | a.ii | a.iii | b.i | b.ii | b.iii | c.i | c.ii | c.iii |
|---|---|---|---|---|---|---|---|---|---|
| $\Delta\overline{P}$ | 10 | 100 | 1000 | 10 | 100 | 1000 | 10 | 100 | 1000 |
| $\overline{\sigma}$ | 1 | 10 | 100 | 0 | 0 | 0 | -1 | -10 | -100 |
| $R_{max}$ (μm) | 11.53 | 11.52 | 11.47 | 11.48 | 11.38 | 11.24 | 11.40 | 11.19 | 10.65 |
| Curvature direction | - | - | - | - | - | - | - | - | - |
| $R_{top}$ $(\mu m)$ | 7.27 | 17.87 | 43.67 | 7.47 | 17.95 | 41.72 | 7.60 | 17.71 | 37.05 |
| $\omega$ $(k_BT/\mu\text{m}^3)$ | 4.00 | 4.63 | 5.35 | 3.12 | 3.17 | 2.97 | 2.32 | 1.72 | 0.09 |
| $\sigma$ $(k_BT/\mu\text{m}^2)$ | 0.37 | 0.61 | 1.02 | 0 | 0 | 0 | -0.34 | -0.62 | -1.42 |
| $\Delta P$ $(k_BT/\mu\text{m}^3)$ | -0.51 | -0.34 | -0.23 | -0.47 | -0.34 | -0.27 | -0.44 | -0.35 | -0.38 |
| $A$ $(\mu\text{m}^2)$ | 1000 | 1000 | 1000 | 1000 | 1000 | 1000 | 1000 | 1000 | 1000 |
| $A_{adh}$ $(\mu\text{m}^2)$ | 313 | 314 | 312 | 298 | 287 | 271 | 279 | 245 | 87 |
| V $(\mu\text{m}^3)$ | 1123 | 1579 | 1724 | 1294 | 1752 | 1956 | 1454 | 1960 | 2482 |
| $\nu$ | 0.38 | 0.53 | 0.58 | 0.44 | 0.59 | 0.66 | 0.49 | 0.66 | 0.83 |
| $A_{adh}/A$ | 0.31 | 0.31 | 0.31 | 0.30 | 0.29 | 0.27 | 0.28 | 0.25 | 0.09 |
